# A genetic model for development, physiology and behavior of zebrafish larvae devoid of catecholamines

**DOI:** 10.1101/2025.07.22.666142

**Authors:** Susana Paredes-Zúñiga, Rebecca Peters, Kristine Østevold, Gerard Arrey, Dennis Frank, Johannes Oswald, Florian Veit, Theresa Schredelseker, Jochen Holzschuh, Wolfgang Driever

**Affiliations:** Developmental Biology, Institute Biology 1, Faculty of Biology, University of Freiburg, Freiburg im Breisgau, Germany; CIBSS and BIOSS - Centres for Biological Signalling Studies, University of Freiburg, Freiburg im Breisgau, Germany

**Keywords:** Catecholamines, dopaminergic neuron, motor behavior, neurogenesis, cardiac physiology

## Abstract

Dopamine and noradrenaline have conserved roles in control of physiology and behaviors of vertebrates. However, vertebrate genetic systems completely devoid of catecholamines are not available. We have generated a genetic zebrafish model completely devoid of catecholamines by combining mutations in all three genes involved in L-DOPA synthesis: the *tyrosine hydroxylase* genes *th* and *th2*, and *tyrosinase tyr*. We found catecholamine-deficient zebrafish larvae to be viable and to develop an anatomically normal nervous system including catecholaminergic neuron somata and projections, albeit with reduced cell numbers detected in some clusters. In contrast, selected physiological functions that depend on catecholamines are impaired, including hatching and heart rate regulation upon temperature challenges. Spontaneous locomotion and optomotor behaviors are also impaired. Despite the changes observed, it is surprising that larvae develop a largely normal behavioral repertoire. Our model will be useful to investigate how physiology and neural circuit function are regulated in catecholamine deficient larvae.

## Introduction

Catecholamines (CAs) are chemical messengers with neuromodulatory and endocrine functions. Endogenous CAs include dopamine (DA), noradrenaline (NA, norepinephrine) and adrenaline (epinephrine)(Moore and Bloom, 1978, 1979; Smeets and Reiner, 1994). Next to the roles of CAs in physiology, progressive loss of DA neurons in Parkinson’s Disease (Hirsch et al., 1988), hypodopaminergia and altered DA-signaling are some of the multiple causes behind neurodevelopmental disorders such as autism (Paval and Miclutia, 2021), and attention deficit / hyperactivity disorder (ADHD) (MacDonald et al., 2024) .

CA neurons can be identified by the expression of the rate-limiting enzyme Tyrosine hydroxylase (TH), which catalyzes the hydroxylation of the amino acid L-tyrosine to L-dihydroxy-phenylalanine (L-DOPA; Supplementary Figure S1A). Therefore, the *Th* gene has become a target for genetic manipulation of CA biosynthesis. The disruption of the *Th* locus in mice leads to mid-gestational lethality(Kobayashi et al., 1995; Zhou et al., 1995). However, surprisingly, *Th* mutant mice have been shown to still contain significant amounts of DA in brain and body(Kobayashi et al., 1995), which has been suggested to derive from maternal bloodstream(Horackova et al., 2022) and from melanocytes(Eisenhofer et al., 2003). In melanocytes, the enzyme Tyrosinase (TYR) catalyzes the oxidation of tyrosine to dopaquinone, a precursor of melanin (Supplementary Figure S1A)(Raper, 1926; Riley, 1999). In redox reactions, dopaquinone is converted to L-DOPA(Eisenhofer et al., 2003), which exits melanocytes and enters the blood stream. In wild type (WT) mice, Tyrosinase does not contribute significantly to DA levels in the brain(Rios et al., 1999). However, in the absence of TH, Tyrosinase-derived L-DOPA partially substitutes NA and DA in peripheral tissues and brain(Rios et al., 1999). For these reasons, genetic mammalian models have so far not been used extensively to address the developmental and behavioral consequences of CA depletion.

Experimental models for DA depletion often use genetic or neurotoxin ablation of DA neurons, and pharmacological or optogenetic manipulation of DA signaling. However, these techniques do not selectively eliminate DA transmission, but also non-DA functions of DA neurons, which typically have dual transmitter phenotypes, either GABAergic or glutamatergic(Borisovska et al., 2013; Chuhma et al., 2009; Descarries et al., 2008; Filippi et al., 2014; Kawano et al., 2006; Vaaga et al., 2014). Further, agonists or antagonist treatments broadly act on CA receptor expressing cells, irrespective of physiological CA signaling, and are prone to cause gain-of-function and potentially indirect effects. Despite the advances in understanding CA systems, we have a limited knowledge of the role of second transmitters in DA neurons, and of the modulatory functions of DA during brain development. Therefore, we aimed at developing a genetic zebrafish model completely devoid of catecholamine transmission, but still containing the cellular complement of wild type CA neurons.

The CA systems in teleosts(Ekström et al., 1994; Meek, 1994), and specifically in zebrafish (Kaslin and Panula, 2001; Ma, 2003; Rink and Wullimann, 2001; Yamamoto et al., 2011), show both conserved and divergent features compared to mammals (Supplementary Figure 1B and legend). While CA systems in the zebrafish adult brain have been extensively described, we will focus here on the experimentally more accessible larval stages (Armbruster et al., 2025; Filippi et al., 2014; Holzschuh et al., 2001; Rink and Wullimann, 2002). In contrast to mammals, zebrafish have two TH encoding genes, *th* (previously also called *th1*), and *th2*(Candy and Collet, 2005; Chen et al., 2009; Filippi et al., 2010; Yamamoto et al., 2010). The *th2* gene likely got secondarily lost in mammals, since it is also found in amphibians and birds(Candy and Collet, 2005; Xavier et al., 2017; Yamamoto et al., 2010). Most *th2* neurons co-express *th*(Fontaine et al., 2015; Semenova et al., 2014) (Figure S1B). The *tyrosinase (tyr)* locus has also been shown to contribute to physiologically relevant biosynthesis of L-DOPA in the retina(Page-McCaw et al., 2004). Therefore, to generate a CA-free model in zebrafish, all three genes, *th*, *th2* and *tyr,* need to be inactivated.

Here, we generated *th*, *th2, tyr* triple mutant zebrafish larvae, validated absence of DA by ELISA, and performed an initial analysis of embryonic and larval development in general and CA systems development specifically. We also characterized potential effects of loss of CAs on cardiac function and basic optomotor behavior. In contrast to CA-deficient mouse models(Kobayashi et al., 1995; Zhou et al., 1995), *th*, *th2, tyr* triple mutant larvae develop largely normal and are viable into larval stages. Mutant larvae however show distinct physiological and behavioral phenotypes affecting hatching, heartbeat regulation and motor behaviors.

## Results

### Triple homozygous *th*, *th2, tyr* mutant larvae are devoid of dopamine

For generating a catecholamine-depleted model, we induced loss-of-function mutations in both *tyrosine hydroxylase* paralogous genes *th* (Figure 1) and *th2* (Figure 2). The *th* gene was mutated using a pair of TALENs targeting the beginning of the ORF in exon 1 containing the start codon ATG and a restriction site for EcoRI (Figure 1A-C). We established a stable mutant line for the allele *th^m1403^* (Figure 1D), which harbors a 7 bp deletion beginning with the G of the start ATG (A equals bp position 1). In the absence of the native TH ATG start codon, the closest ATG is located out-of-frame at 125 bp, followed by a stop codon after 18 aa (Figure 1E). Anti-TH immunohistochemistry with a polyclonal zebrafish TH antibody raised against amino acids 92 to 365(Kastenhuber et al., 2010) shows the complete absence of TH immunoreactive proteins in *th^m1403/m1403^* larvae (Figure 1F), demonstrating that the *m1403* allele is a protein Null allele.

**Figure 1.**
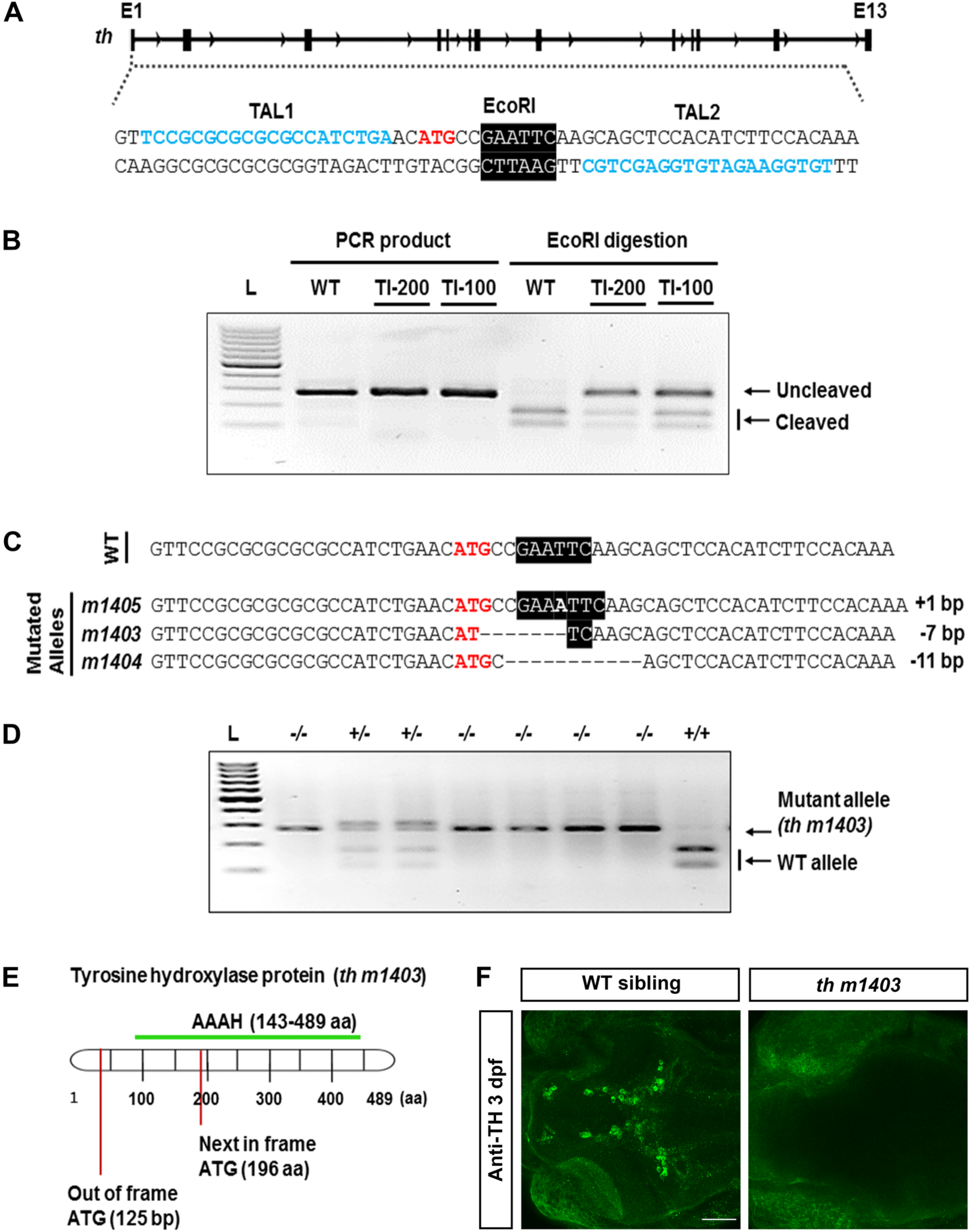
TALEN-mediated mutation of the *th* locus. A) TALEN were designed to prevent Th translation by disrupting the beginning of the ORF. Schematic representation showing the binding sites for TALEN pairs (TAL1 and TAL2, blue) in the exon 1 (E1). The start codon ATG is shown in red and the recognition site for EcoRI is highlighted in black. B) TALEN efficacy assay of injected G0 embryos. PCR amplification followed by EcoRI digestion shows that TALEN-injected embryos present an uncleaved band, indicating an indel mutation at the target site. TALEN encoding mRNAs were injected at two different concentrations 100 ng/µl (TI-100) and 200 ng/µl (TI-200). WT: wild type, L: 100 bp ladder. (C) Sequencing results of the undigested bands from individual F1 heterozygous fish. On top, reference wild type allele (WT). Mutated alleles: *m1405* (+1 bp addition), *m1403* (-7 bp deletion), *m1404* (-11 bp deletion). (D) *th^m1403^* genotyping assay reveals 278 bp fragment in mutant (-/-) and EcoRI cleaved 167 and 111 bp fragments in WT (+/+). (E) Scheme showing the next in and out of frame ATGs after the start-ATG in the Th protein. The succeeding in-frame ATG, which could serve as an alternative initiation codon, is at 589 bp (ORF amino acid 196) at the beginning of the Th active aromatic amino acid hydroxylase domain. Translation from this start codon would lead to a hypothetical carboxyterminal protein of about 60% of the Th sequence. AAAH, green: active aromatic amino acid hydroxylase domain, L: 100 bp ladder. (F) Anti-Tyrosine hydroxylase immunofluorescence in 3 dpf WT siblings and homozygous *th* mutants show complete absence of Th immunoreactivity in *th^m1403/m1403^* larvae. Dorsal view, anterior left. Scale bar: 50 µm.

**Figure 2.**
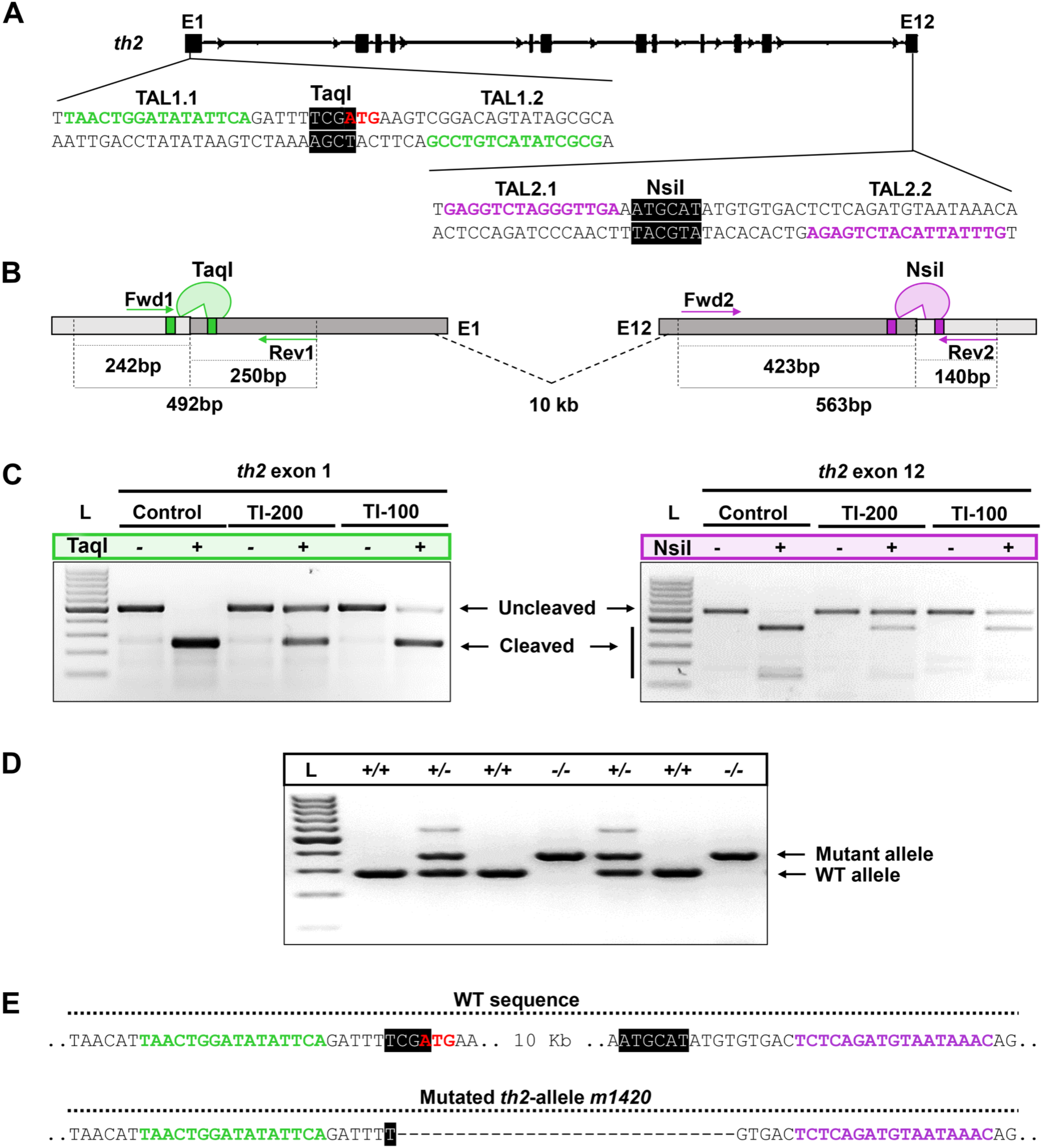
TALEN-mediated strategy for *th2* deletion. TALENs were designed for targeting the *th2-*exon1 (E1) and *th2*-exon12 (E12). Black boxes depict the 12 *th2* exons. Close-up to the TALEN-binding regions in exons E1 (TAL1.1 and TAL1.2, green) and E12 (TAL2.1 and TAL2.2, magenta). ATG start codon is shown in red, TaqI (*th2*-E1) and NsiI (*th2*-E12) sites are highlighted. (B) Scheme of restriction fragment length polymorphism (RFLP) strategy for genotyping and expected PCR product sizes; bp: base pairs. (C) Digested PCR products amplified from *th2*-E1 and *th2*-E12. Control: non-injected samples; TI-200: TALEN-injected samples with 200 ng/μL; TI-100: TALEN-injected samples with 100 ng/μL; (+) restriction digest; (-) non-restricted; L: 100 bp ladder. (D) Complete deletion of *th2* E1-E12 in the *th2^m1420^* allele. For *th2^m1420^* genotyping, a common reverse primer (5’-ATGCCCCTGGAAATAGGC-3’) and two mismatch forward primers hybridizing either the wild type (5’-CTTGGGGAGACTGGGACAG-3’) or mutant (5’-CTGTCGCTTGAAGCACCTG-3’) sequences were used, resulting in DNA fragments of 286 bp for wildtype (+/+) and 396 bp for mutant alleles (-/-); L: 100 bp ladder. (E) DNA sequences from control (WT sequence) and *th2* deletion (mutated *th2*-allele *m1420*).

To ensure the complete knock-out of *th2*, a deletion of all coding exons was performed. (Figure 2). TALEN-binding regions included a restriction site for TaqI in *th2*-Exon1 and NsiI in *th2*-Exon12 (Figure 2A-C). The 10 kb deletion allele *th2^m1420^* was confirmed by allele-specific PCRs across the deleted region and sequencing of the Exon1-Exon12 linking fragment (Figure 2D, E).

We determined DA content of WT and mutant larval heads using an ELISA test. Given that commercial kits are not designed for zebrafish extract samples, we first determined whether there is a linear relationship between the amount of extract from 5 dpf larvae and the ELISA DA readings. A two-step serial dilution of WT extracts reveals a near-linear relation between the sample amount in the assay and the DA amounts calculated based on ELISA readings (Figure 3A). As negative control we used 20 hpf embryo extracts, which should be devoid of DA because *th*, *th2* and *tyr* expression have not started at this developmental stage (Figure 3A), and indeed did not detect DA above background.

**Figure 3.**
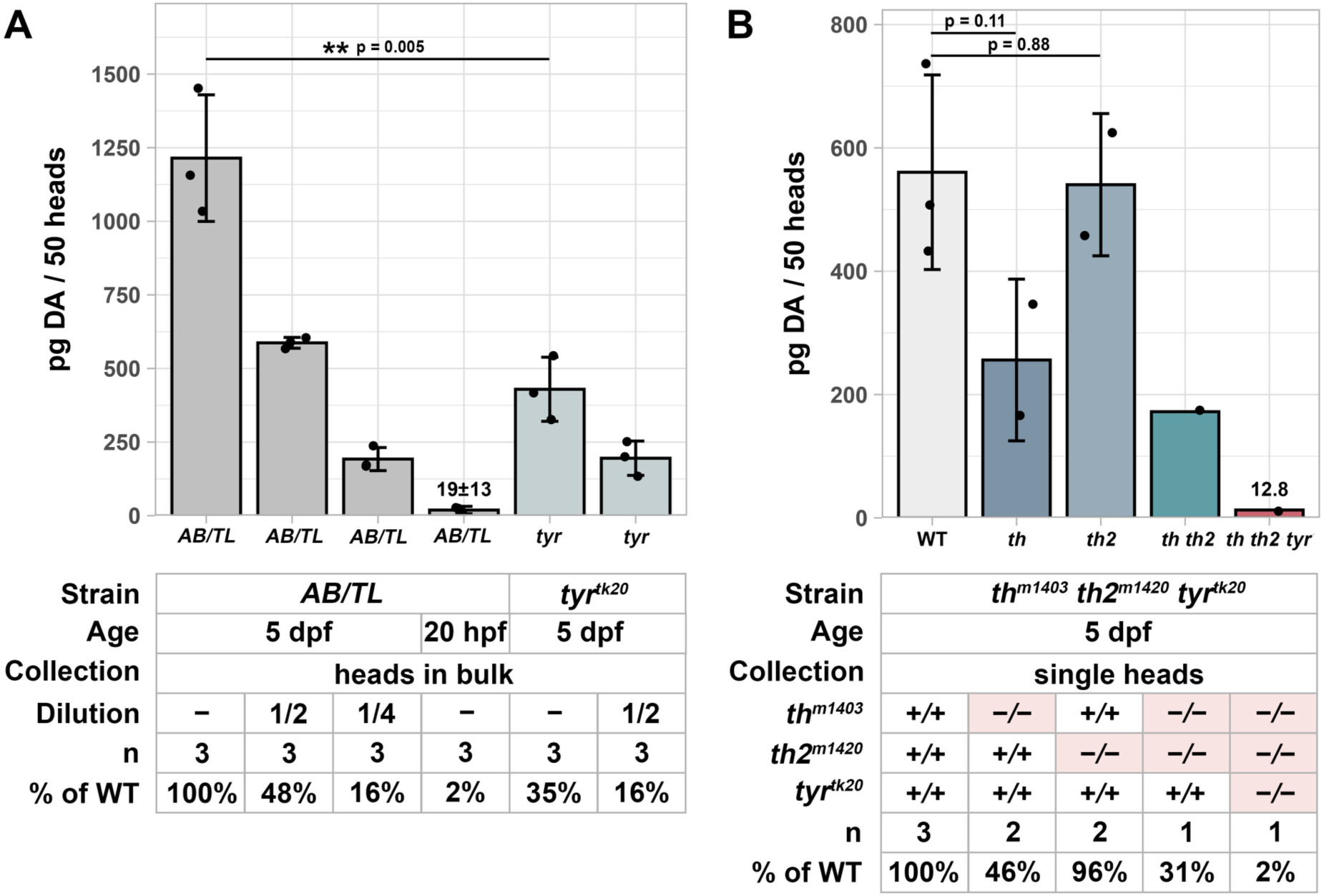
Dopamine depletion in *th, th2, tyr* mutant embryos. (A) Dopamine amount detected by ELISA in 50 bulked heads. From left: Sample dilution series of WT AB/TL samples show a linear ELISA readout. Measurements were obtained from 500, 250 or 125 µl extracts obtained from 50 bulked heads of 5 dpf larvae (see methods). As negative control 20 hpf AB/TL whole larvae were used, as at this stage Tyr or TH activities are still not expressed. Measured DA concentration from 500 or 250 µl extract from 50 *tyr*-/-bulked heads of 5 dpf larvae. (B) Dopamine amounts measured in 50 bulked heads of single, double and triple mutant *th, th2,* and *tyr* genotypes. Sample collection was done at 5 dpf using heads pooled after genotyping of trunk material from individual heads (single heads). Samples were applied in biological triplicates or duplicates, if feasible. Statistical analysis was performed using one-way ANOVA (see Supplementary Table S1).

To determine relative contributions of each gene to total DA, we performed ELISA on wild type (WT) and single, double, and triple *th, th2, tyr* mutant combinations. The results indicate that in *th* (Figure 3B; 46% DA of WT; p=0.11) and *tyr* (Figure 3A; 35% of WT; p=0.005) mutants DA levels are reduced compared to WT. In *th2* mutants (95% DA of WT) the observed reduction is not significant (Figure 3B; p=0.88). *th, th2* double mutants are not sufficient to completely impair DA biosynthesis (Figure 3B), as 31% of WT DA levels remained in the double mutants. In contrast, *th, th2, tyr* triple mutants reveal absence of DA, as the measured DA amount is not different from 20 hpf WT negative control samples (Figure 3A, B). Since DA is the first CA synthetized from L-DOPA, absence of DA indicates that triple mutants are also devoid of NA and adrenaline.

### Catecholaminergic cluster development in *th th2* double mutant larvae

To investigate a potential impact of absence of *th* and *th2* activity on DA neuron development, we analyzed *th* expression by whole mount in situ hybridization (WISH). Since the *th^m1403^* allele impairs TH translation but not *th* transcription, *th* WISH labels DA neurons in *th th2* double mutant larvae. While in *th th2* mutant larvae CA neurons were detected in all anatomical clusters at 4 dpf, compared to WT, a reduced *th* WISH stain intensity was observed in several clusters, including hindbrain locus coeruleus (LC) and the telencephalic olfactory bulb (OB) and subpallial (SP) groups (Figure 4A). This difference may be caused by *th^m1403^* transcripts being potentially less stable due to nonsense-mediated mRNA decay (Brogna and Wen, 2009; Wittkopp et al., 2009). We counted *th* WISH positive cells in *th th2* double mutants and WT siblings (Figure 4C, Supplementary Table S4). DA neurons of the OB and SP, as well as of the diencephalospinal DA system clusters DC4 and DC5, were counted combined, as the anatomical borders between each pair of clusters could not be determined in the image stacks. Hypothalamic DC7 neurons were not counted, because both in WT and mutants the stain intensity was too low to allow reliable cell counting. We detected a significantly smaller number of DA neurons in prethalamic DC1 and preoptic (PO) DA clusters (Figure 4A,C). However, even in WT larvae PO and DC1 DA neurons express low levels of *th*, so that *th* expression in *th th2* mutants in some DA neurons may have been below the detection limit.

**Figure 4.**
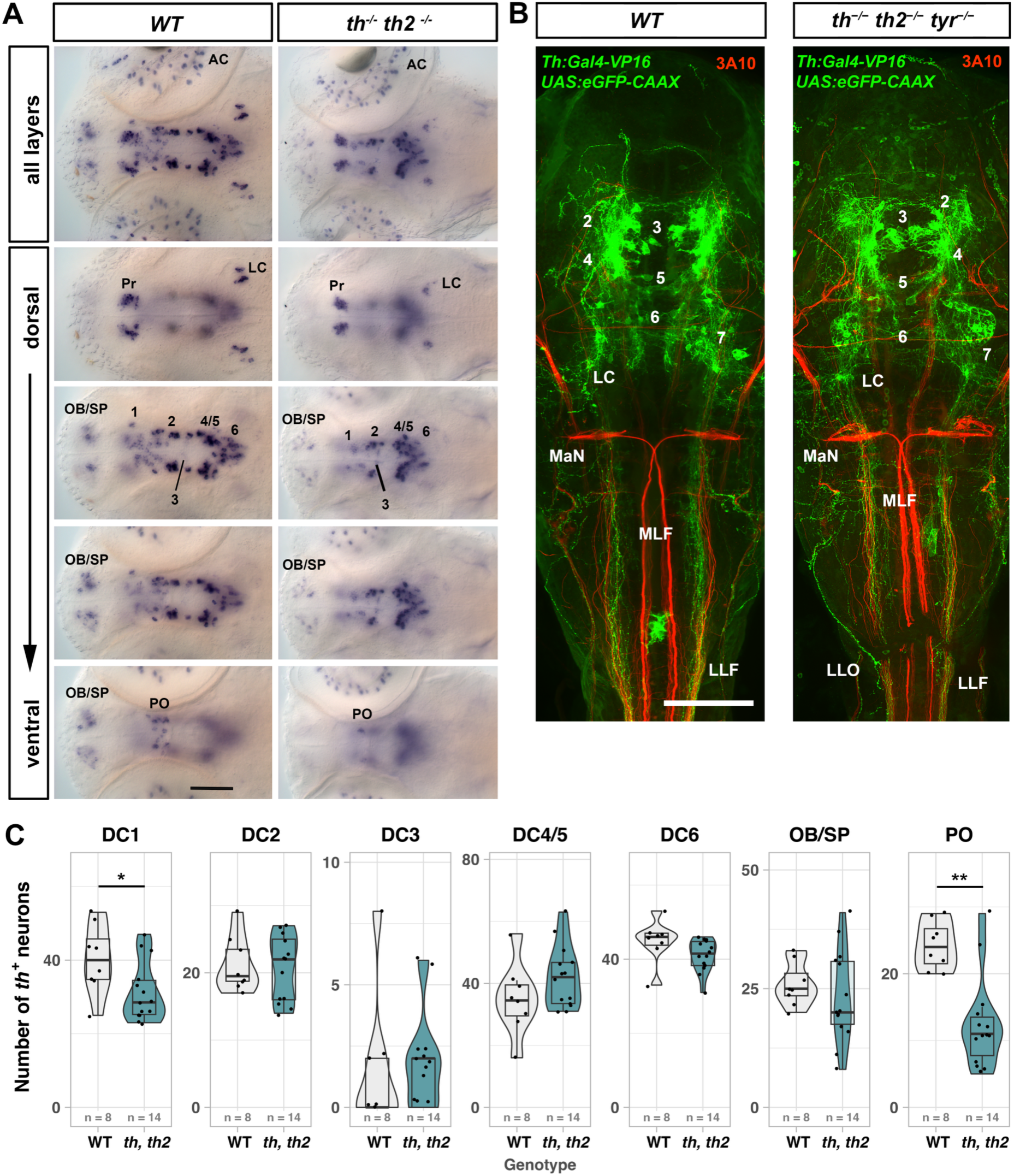
Development of dopaminergic neurons and their projections in *th, th2* deficient larvae. **(A)** WISH showing *th* expression in 4 dpf WT (n = 8) and *th, th2* double mutant (n = 14) larvae. Images show z-projections of DIC optics stacks with 1 µm distance between optical planes; dorsal views, anterior left. Optical planes (#1 is most dorsal) used for Z-projections are from top to bottom: WT all 1-108, dorsal 1-31, 45-60, 56-83, ventral 78-96; *th th2* double mutants all 1-100, dorsal 23-35, 56-63, 52-65, ventral 78-88. Scale bar: 100 µm. **(B)** Anti-GFP (green) and anti-3A10 (red) immunofluorescence staining in 3 dpf *Tg(th:Gal4-VP16)^m1233^, Tg(UAS:eGFPCAAX)^m1230^* transgenic larvae show CA tracts and axonal projections (green) and other axonal scaffolds (red). Genotypes as indicated (n = 2 for each genotype), dorsal view, anterior up. Scale bar: 100 µm. (C) Counts of *th*-expressing cells present in the dopaminergic clusters of *th, th2* double mutants and WT siblings shown in (A). A significant reduction of DA neurons was detected for PO (p-value = 0.002) and DC1 (p-value = 0.04) groups (unpaired two-tailed Mann-Whitney test, Supplementary Table S2). Abbreviations: 1-7 or DC1-7 Ventral forebrain DA groups 1 to 7; AC, Amacrine Cell; DC, diencephalic groups; LC, locus coeruleus; LLF, lateral longitudinal fascicle; LLO, lateral line organ; MaN, Mauthner neuron; MLF, medial longitudinal fascicle; MO, medulla oblongata; OB, olfactory bulb; PO, preoptic region; Pr, pretectum; SP, subpallium.

To determine whether CA deficiency affects axonogenesis of CA neurons, we crossed a 3 dpf transgenic line expressing membrane-anchored GFP in *th* cells (*Tg(th:Gal4-VP16)^m1233^*, *Tg(UAS:eGFP-CAAX)^m1230^*) into a *th, th2, tyr* heterozygous line. We combined anti-GFP immunofluorescence for CA axons with anti-neurofilament 3A10 stain for the general axonal scaffold. Staining of axonal projections shows no major differences between WT and CA-deficient larvae (Figure 4B). In triple mutants, axons of the medial longitudinal catecholaminergic tracts project caudally from the posterior tuberculum through midbrain and hindbrain just medial to the lateral longitudinal fascicle into the spinal cord. The lateral line and other peripheral sense organs also show normal innervation from catecholaminergic tracts(Haehnel-Taguchi et al., 2018). The triple mutants also reveal that in CA-deficient larvae, similar to *th th2* double mutants, all anatomical CA groups develop. Our finding that GFP expression from the *th:Gal4* BAC transgene driven by *th* genomic control region appears at similar levels in WT and *th, th2, tyr* mutants (Figure 4B) supports our notion that expression from the *th* promoter may indeed not be affected in CA-deficient larvae.

A similar analysis of *th2* expression in *th2* DA neurons was not possible because the *th2* mutant allele deletes most of the *th2* transcribed region. However, given that many DC7 DA neurons express both *th* and *th2* (Chen et al., 2009; Filippi et al., 2010; Yamamoto et al., 2010), and that DC7 neurons are present in triple mutants (Figure 4B), we postulate that *th2-*type DA neurons also develop in the triple mutants.

### Spinal motor neuron numbers are not affected by catecholamine deficiency

DC2 and DC4 A11-type DA neurons with descending diencephalospinal projections have been reported to promote motor neuron (MN) generation in the zebrafish spinal cord (Reimer et al., 2013). These experiments used Morpholino knockdown of *th* and DA signaling pathway components, as well as pharmacological DA antagonists, and therefore DA was not completely eliminated. We tested whether complete elimination of DA affects spinal MN development. We crossed *Tg(mnx1:GFP)^ml2Tg^*, which labels primary and secondary MNs, into *th, th2, tyr* line, performed anti-GFP immunofluorescence on embryos fixed at 33 hpf or 3 dpf (Figure 5A), and counted labelled MNs in spinal sectors correlating in length to one somite. *mnx1:GFP* brightly labels new primary and secondary MNs, but expression decreases in older MNs, such that MN counts were lower at 3 dpf. The relative immunofluorescence intensities were not different between mutants and WT. No significant differences were detected when counts of MNs in WT, *tyr* and triple mutant genotypes were compared, neither in the number nor location of MNs (Figure 5B, Supplementary Tables S3, S4). Therefore, we did not observe an effect of DA deficiency on MN development, suggesting that the previously reported requirement for DA(Reimer et al., 2013) may actually have been caused by experimental interference with other non-DA regulatory systems.

**Figure 5.**
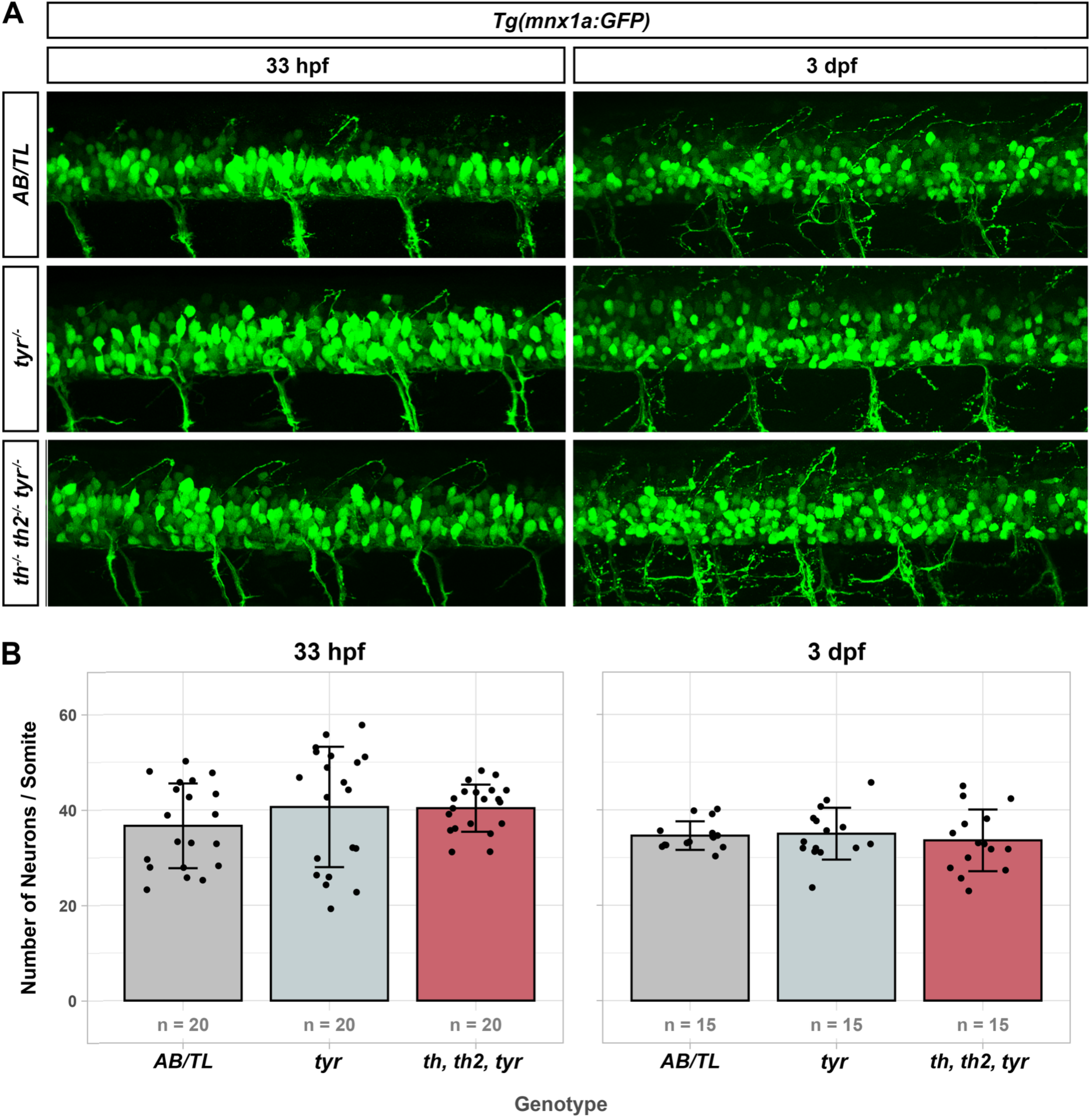
Primary and secondary motor neuron numbers are not affected by catecholamine depletion. (A) GFP immunofluorescence in *Tg(mnx1a:GFP)^ml2Tg^*embryos show no difference between the genotypes examined, neither at 33 hpf nor at 3 dpf. Lateral view, anterior left, dorsal at top. (B) *mnx1a:*GFP positive primary (PMN) and secondary (SMN) motor neurons were counted at 33 hpf in 4 somites (7, 8, 9 and 10) of 5 embryos each, and at 3 dpf in 3 somites (7, 8 and 9) of 5 embryos each. At 3 dpf *Tg(mnx1a:GFP*) expression is low in some cells, which were not counted, resulting in smaller cell numbers counted at 3 dpf. For PMN: *F* Welch (2, 33.3) = 1.37, *p* = 0.27. For SMN: *F* Welch (2, 24.99) = 0.21, *p* = 0.81 (Supplementary table S3, S4). Mean with SD are plotted.

### Catecholamines are necessary for HGC granule secretion

At a gross anatomical level, CA-free larvae appear to develop largely normal up to 5 dpf (Figure 4B). However, a careful inspection revealed major differences in hatching gland cell (HGC) retention during larval stages (Figure 6A). The secretory hatching gland epithelium produces and releases proteolytic enzymes involved in the breakdown of the chorion during hatching(De la Paz et al., 2017), and HGCs disappear following this process. Therefore, most HGCs are lost in larvae older than 2 dpf: 200 WT larvae analyzed at 3 dpf were all completely devoid of HGCs. In contrast, in *tyr-/-* larvae we found 9.5% HGC retention at 3 dpf (n=16 out of 168), and 2.9% HGC retention at 5dpf (n=5 out of 168). Thus, in *tyr-/-* mutants a small but significant portion of larvae retains HGCs to 5 dpf. To efficiently recover triple mutants for our analysis, we used a *th+/-, th2+/-, tyr-/-* incross, with all progeny *tyr* mutant. We observed that 24.7% of the progeny from this cross (n=39/158) had HGCs still detectable at 3dpf (HGC[+], Figure 6A). Genotyping revealed a correlation between HGC retention phenotype and *th* genotype (Figure 6B). 68.3% of the HGC[+] larvae were *th* mutants (*th, tyr* double or *th, th2, tyr* triple mutants), while only 6.4% of all HGC[-] larvae were *th* mutants. Accordingly, 93.6% of the HGC[-] larvae had at least one WT *th* allele (Figure 6B). These results show that animals lacking TH have impaired mechanisms of HGC granule secretion, demonstrating that *th* gene activity is necessary for the normal function of HGCs. We analyzed expression of the HGC specific marker *cathepsin Lb* (*ctslb)*(Thisse et al., 1994) and found expression maintained in HGC[+] larvae at 4 dpf (Figure 6C). When we reexamined 3 dpf HGC[+] larvae at 5dpf (Figure 6D), we observed three different phenotypes in these larvae: retention of all HGCs (High), few HGCs (Low) and complete absence of HGCs (None, Figure 6D), revealing progressive loss of HGCs during larval stages in some genotypes (Figure 6E). After genotyping all larvae, we noticed that high retention at 5 dpf was only observed in *th-/-* mutants (Figure 6E, Supplementary Table S6). However, a fraction of *tyr* single mutant larvae also shows low HGC retention. Tyrosinase derived L-DOPA more likely contributes to tissue level interstitial DA, rather than affecting synaptic transmission. We hypothesize therefore that tissue CA levels may directly affect HGC secretion, potentially independent of the established DA-prolactin neurosecretory hypothalamic-hypophyseal axis(De la Paz et al., 2017; Schoots et al., 1982; Schoots et al., 1983). To elucidate whether DA or NA receptors on HGCs may be able to mediate direct effects of systemic CA levels, we queried the Daniocell database (www.daniocell.nichd.nih.gov) (Farrell et al., 2018) for expression of CA receptors in *ctsl1b* expressing HGC clusters (clusters axia.15, axia.18, axia.20; Supplementary Figure S2A). The results indicate that HGCs express *drd5a* and *adrb3a* at higher levels, as well as *adrb1, adrb2a* at lower levels, which are all adenylate cyclase activating receptors (Supplementary Figure S2B-G). Thus, DA or NA may potentially stimulate HGC secretion directly.

**Figure 6:**
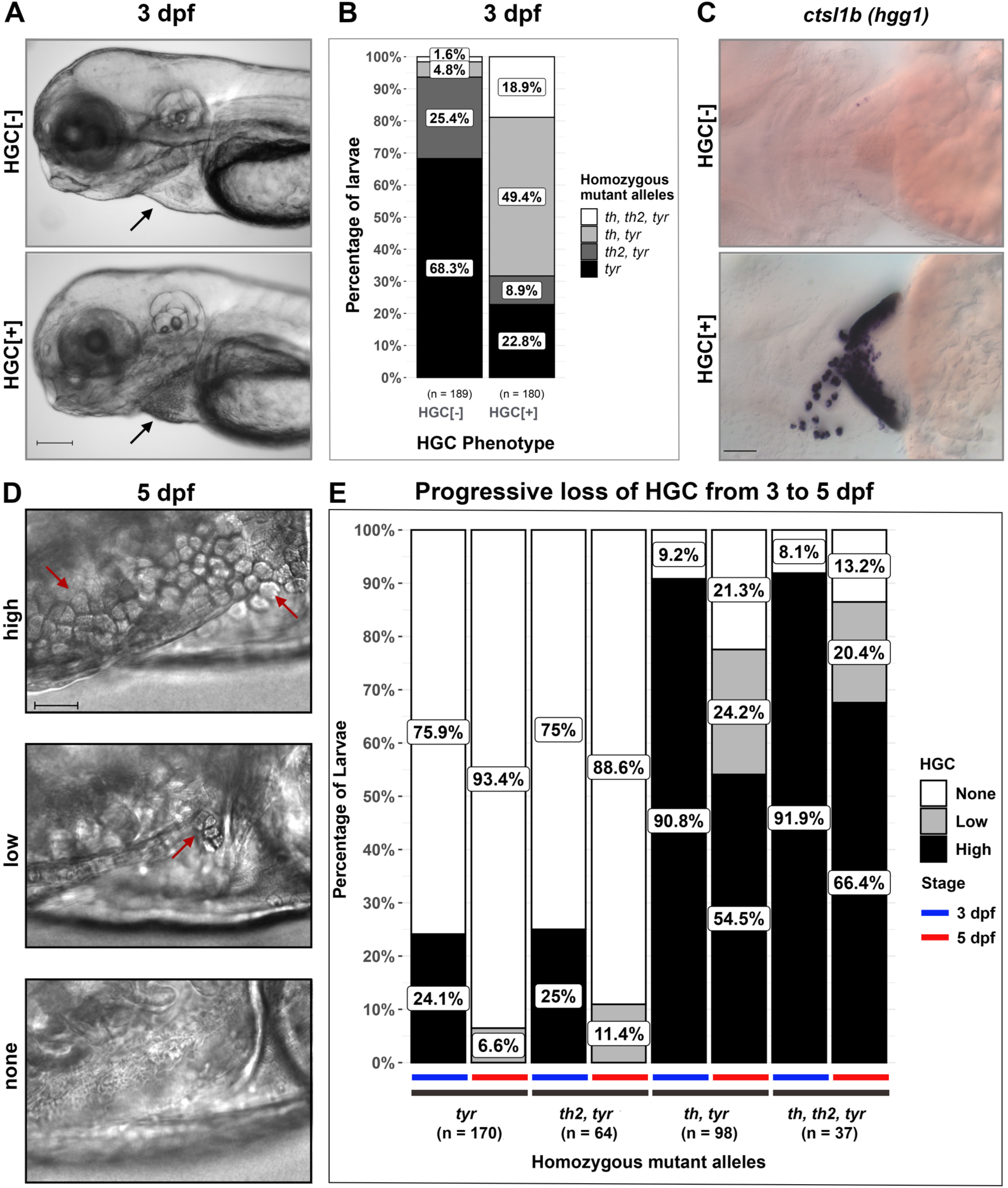
HGC retention is a characteristic phenotype of *th* mutant larvae. (A) HGC phenotype at 3 dpf larvae obtained from a *th+/-, th2+/-, tyr-/-* cross, live images, lateral view, anterior left. HGC[-]: absence of granule-containing cells in *th +/-, th2 +/+, tyr-/-* larva; HGC[+]: retention of granule-containing cells in triple mutant larva. Black arrow points towards the pericardial region where HGCs are located. Scale bar 100 μm. (B) Percentages of larval *th* and *th2* genotypes found in HGC phenotypes of *tyr*-/- larvae (for *tyr* mutants, *th* and/or *th2* may be heterozygous; for *th, tyr* mutants, *th2* may be heterozygous; for *th2*, *tyr* mutants, *th* may be heterozygous). The HGC-retention phenotype correlates with the *th* genotype (Pearson’s chi-square χ^2^ (3) = 152.7, p-value < 6.88e-33, *V* Cramer = 0.64, CI 95% [0.55, 1.00]). (C) WISH showing *ctslb* (*cathepsin Lb;* previously *hgg1*) expression in larvae with either HGC[-] in *th+/+, th2+/+, tyr-/-* larva or HGC[+] in triple mutant larva at 4 dpf. Scale bar 40 μm. (D) Live images of hatching gland region in 5 dpf larvae from a *th+/-, th2+/-, tyr-/-* cross (anterior left, dorsal up). Larvae classified as HGC[+] at 3 dpf, when reanalyzed at 5 dpf, reveal progressive HGC loss during larval development. Red arrows point at exemplary HGCs. High, low and none describe the different degrees of HGC retention observed. Genotypes shown are “High” HGC retention: *th-/-, th2+/-, tyr-/-.* “Low”: *th+/+, th2+/+, tyr-/-.* “None”: *th+/-, th2+/-, tyr-/-*. Scale bar 20 μm. (E) HGC retention in larvae obtained from a *th+/-, th2+/-, tyr-/-* cross were analyzed at 3 dpf and again at 5 dpf. The graph shows percentage of larvae classified as high, low or no HGC retention for each genotype at 3 dpf (left bar, blue line) and 5 dpf (right bar, red line); for genotypes see explanation in (B).

### Catecholamines regulate heart rate and heart rate variability upon thermal challenge

In response to acute physiological stress, the mammalian sympathetic nervous system triggers a multisystemic CA reaction responsible for the physiological changes needed to cope with sudden environmental threats(Seebacher, 2009). Sympathetic modulation of heart performance involving CAs is key for a fast “fight or flight” response. Since our model is completely devoid of CAs, we investigated how CA deficiency affects basic cardiac parameters, such as heart rate (HR) and heart rate variability (HRV), during acute variation of temperature conditions(Barrionuevo and Burggren, 1999; Gierten et al., 2020). Heartbeat was recorded in 3 dpf non-anaesthetized and freely moving larvae at control temperature (28°C) and at four thermal challenges (20°C, 24°C, 32°C and 36°C), each measured 1 min after sudden temperature shift. At 28°C, we observed in *tyr-/-* larvae a HR of 191 beats per minute (bpm; n=18, STDEV 9.9 bpm), similar to previous reports for wild type larvae at this developmental stage(Joyce et al., 2022). HRV was 35.95 ms (n=18, STDEV 7.14 ms), also similar to previously reported values (30 ms)(Harrington et al., 2017). As expected from an ectotherm animal, we observed a positive correlation between temperature and HR (Supplementary Figure S3A). Considering all temperature conditions and all genotypes, two-way ANOVA revealed a significant interaction between temperature and genotype for HR and HRV (HR *p* = 0.00386, HRV *p* = 0.00078; Supplementary Table S8A and S8B), indicating that both HR and HRV responses to temperature changes depend on the genotype of the fish. To specifically identify which genotypes contribute to these differences, Tukey’s test was conducted and revealed that *th*, *th2*, *tyr* triple mutants exhibited significant differences in HR and HRV when compared to *tyr* single mutants across specific temperature conditions (Tukey’s multiple-pairwise comparisons, HR *p* = 0.0259, HRV *p* = 0.00011, Supplementary Table S8C and S8D terms 11-16). When analysing specific temperatures, no differences in HR or HRV were detected when mutant genotypes were compared at standard breeding temperature (28°C) (Figure 7A, B and Supplementary Table S8C and S8D terms 17-206 rows highlighted yellow). However, for HR, at 36°C *th, tyr* double mutants show a significantly smaller HR than *tyr* mutant or *th2, tyr* mutant siblings (Supplementary Table S8C rows highlighted green). For HRV, both *th, tyr* double and *th, th2, tyr* triple mutants have a significantly smaller HRV at 24°C and 20°C when compared to *tyr* single mutants (Figure 7B, Supplementary Table S8D rows highlighted blue). Our data thus reveal that in zebrafish larvae CAs contribute to temperature dependent HR and HRV modulation.

**Figure 7.**
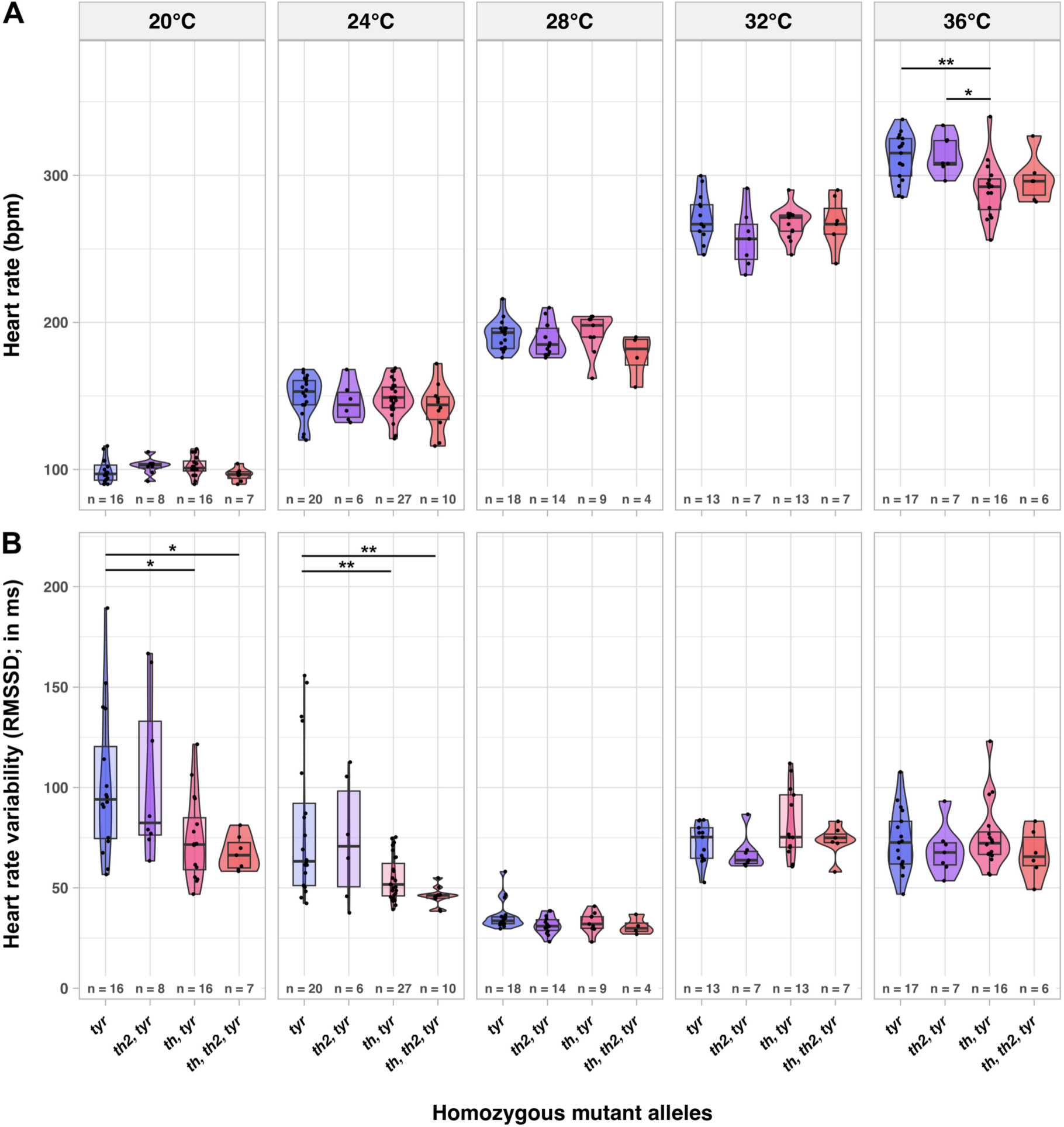
Reduced catecholamine levels affect normal HR and HRV adaptations during a thermal challenge. (A) Heart rate (HR) and (B) heart rate variability (HRV) for *th* and *th2* genotypes in a *tyr* mutant background (3 dpf) at different temperatures. Statistical analysis was performed using two-way ANOVA type III with Tukey’s multiple-pairwise comparisons test (See supplementary Table S8). Only significant differences are shown.

At standard breeding temperature (28°C) no differences in HR or HRV were detected when mutant genotypes were compared (Figure 7A,B and Supplementary Table S8). However, comparing all genotypes at all temperatures combined reveals a significantly reduced HR of *th, th2, tyr* triple compared to *tyr* single mutants (Tukey’s multiple-pairwise comparisons, *p* = 0.0259, Supplementary Table S8). Two-way ANOVA further revealed a significant interaction between temperature and genotype for HR and HRV (HR *p* = 0.00386, HRV *p* = 0.00078; Supplementary Table S8). For HR, only at 36°C *th, tyr* double mutants show a significantly smaller HR than *tyr* mutant or *th2, tyr* mutants siblings. For HRV, both *th, tyr* double and *th, th2, tyr* triple mutants have a significantly smaller HRV at 24°C and 20°C when compared to *tyr* single mutants (Figure 7B, Supplementary Table S8). Our data thus reveal that in zebrafish larvae CAs contribute to temperature dependent HR and HRV modulation.

### Catecholamines modulate spontaneous and stimulus-triggered movement

DA plays a crucial role in zebrafish normal locomotor development(Lambert et al., 2012; McPherson et al., 2016) as well as in sensory processing(Mu et al., 2012). Therefore, we assessed if CA depletion specifically affects locomotor activity. To avoid effects of genetic heterogeneity in the non-inbred zebrafish strains, in each experiment we used larvae from a single *th+/-*, *th2+/-, tyr-/-* cross. To recover sufficient numbers of triple mutant embryos from an individual cross, we used *tyr-/-* larvae as internal reference instead of WT larvae. *tyr* mutant and WT AB/TL larvae have similar spontaneous motor activity based on analysis of distance traveled (Supplementary Figure S4).

We first determined spontaneous movement measured as distance swam in 1 hour by each larva in a bright environment without specific stimuli. We observed that mutant combinations including *th* (*th-/-, tyr-/-* and *th-/-, th2-/-, tyr-/-*) show a significantly reduced spontaneous locomotion compared to *tyr* siblings (Figure 8A). In contrast, *th2 tyr* mutants showed no significant locomotor activity differences when compared with *tyr* siblings (Figure 8A). These results indicate that CA modulation of spontaneous swimming depends predominantly on TH but not on TH2 activity. Next, we examined whether a visual stimulus triggered response was also affected by CA depletion. We analyzed the optomotor response (OMR), a reflexive behavior where the animal follows the direction of whole-field motion(Kist and Portugues, 2019). Moving gratings were presented to individual larvae from below, orthogonal to the longitudinal swim rails, and larvae recorded (Figure 8B). We classified the motor outputs in three groups: OMR (moving in the direction of the stripes), no-OMR (moving without engaging in OMR) and no-movement (remaining still during the entire trial; Figure 8C). For all genotypes the vast majority of larvae engages in OMR . However, we observed small differences in the percentages of animals that do not engage in OMR or do not move. Surprisingly, for all genotypes that contain *th2-/-* mutants, larvae appear to have slightly higher OMR response percentages, which may relate to *th2* being mostly expressed in the hypothalamus, where basic behavioral states including sleep and alertness are regulated. However, the relatively small differences may not justify a mechanistic interpretation.

**Figure 8.**
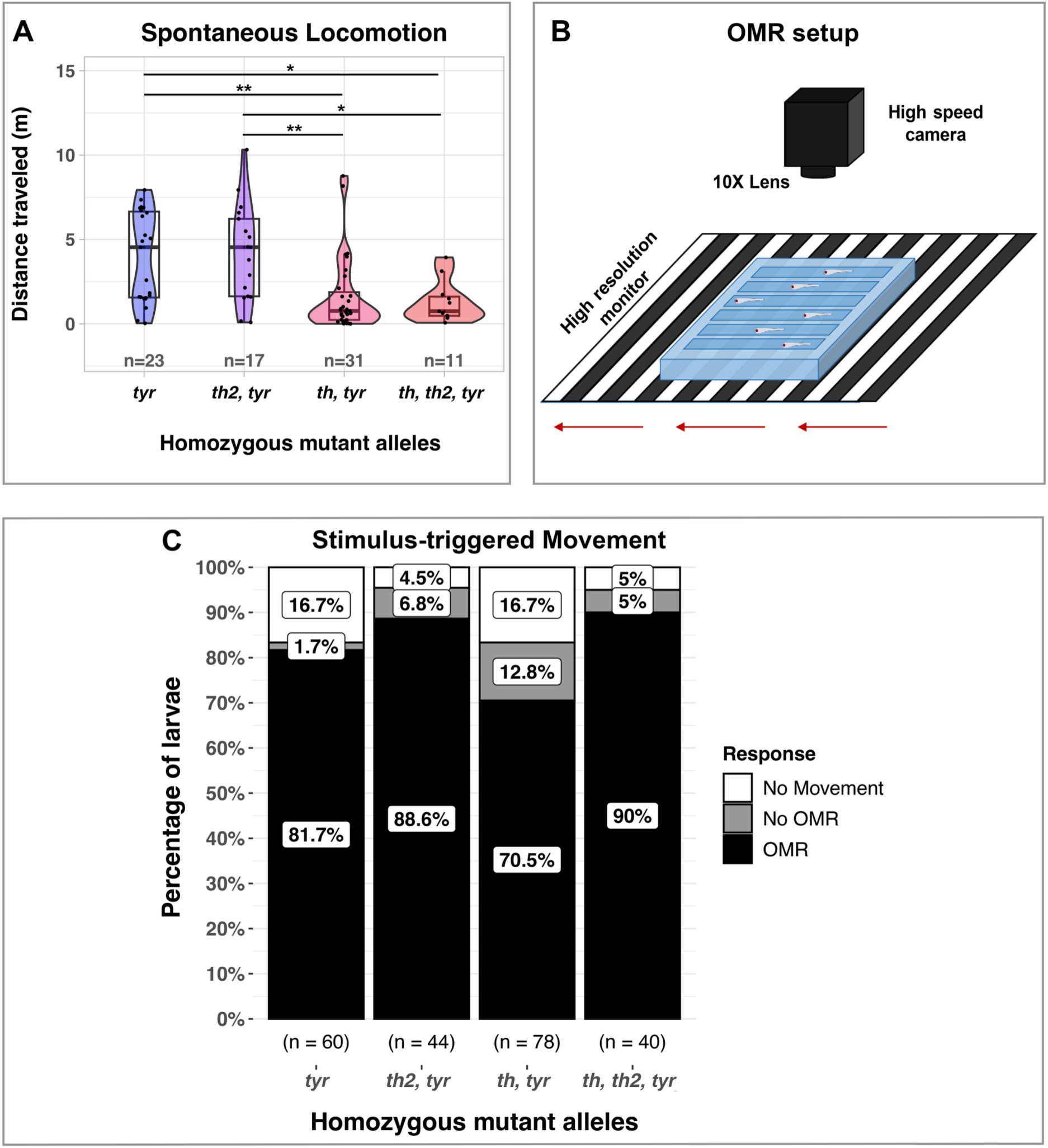
*th* derived catecholaminergic activity is required for normal locomotor activity. (A) Spontaneous locomotor activity measured as distance traveled in 1 hour (hr) is reduced in 5 dpf *th* mutant compared to *th2* mutant or *tyr* control larvae. Statistical analysis was performed using Kruskal-Wallis rank sum test (χ2 (3) = 19.95, *p* = 0.000174, effect size (ε^2^ (rank) = 0.25, CI 95% [0.12, 1.00]), with Dunn’s pairwise comparisons test. *: p-value = 0.03; **: p-value = 0.003. (B) Schematic representation of optomotor response (OMR) setup. Red arrows indicate grating movement. (C) Comparison of response types reveals that all genotypes perform OMR with minor differences in efficiency (Pearson’s chi-square χ2 (6) = 14.117, p-value < 0.028, *V* Cramer = 0.14, CI 95% [0.00, 1.00]). See also Supplementary Figure S5 for outlier analysis.

Since OMR depends on both visual and motor systems function, DA impairment may differentially affect the kinematic parameters composing the OMR. We measured the trajectory and displacement as well as the time taken by the larvae to reach the end of the OMR track. The latency in responding to the stimulus onset was also determined. *th, th2, tyr* mutants have a higher latency in initiating an optic-flow directed movement than *tyr* or *th2, tyr* mutant siblings (Figure 9A). These data reveal that OMR latency is increased by the lack of TH and TH2 activity. Based on trajectory, displacement and time taken, we calculated directionality (Figure 9B), velocity (Figure 9C) and speed (Figure 9D). Both velocity and speed are significantly reduced in *th* mutants (*th, tyr* and *th, th2, tyr*) when compared with the *tyr* control or *th2, tyr* mutant siblings. Upon excluding outliers, also directionality appears significantly reduced (Supplementary Figure S5B). Lack of *th2* activity does not affect any of the four parameters significantly. While latency and directionality are related to proper detection and processing of visual cues, the speed and velocity largely depend on downstream motor pathways, suggesting that CA modulation affects both visual and motor components of the OMR.

**Figure 9.**
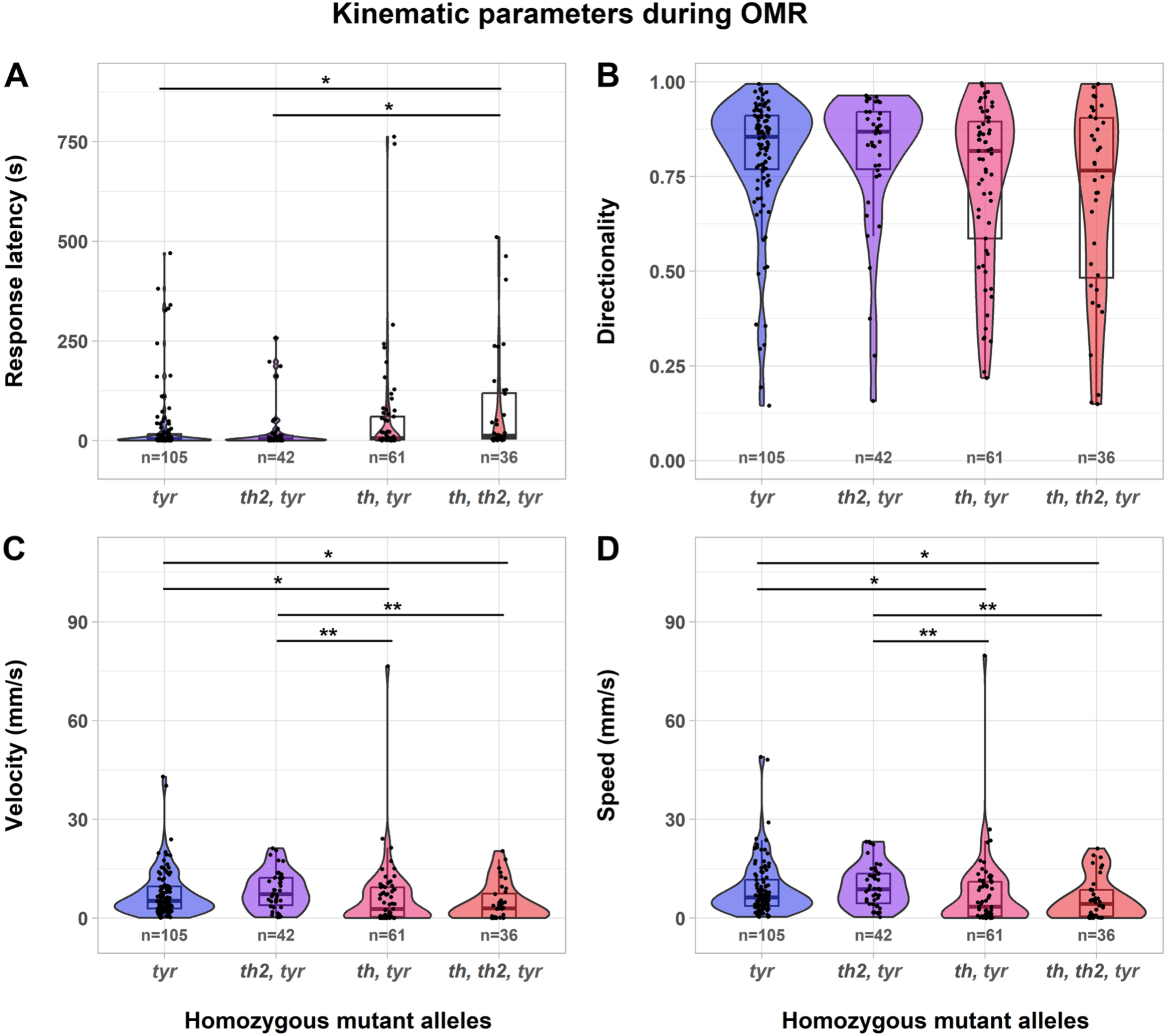
Catecholamine deficiency affects OMR kinematic parameters. Kinematic parameters during optomotor response (OMR) for each larval genotype at 5 dpf. (A) Latency in OMR initiation: Delay in responding to the stimulus onset (χ2 Kruskal-Wallis (3) = 14.02, *p* = 0.00288, effect size (ε^2^) = 0.06, CI 95% [0.03, 1.00]). (B) Directionality: Quotient between displacement and trajectory (χ2 Kruskal-Wallis (3) = 7.73, *p* = 0.05, effect size (ε^2^) = 0.03, CI 95% [0.01, 1.00]). (C) Velocity (vector): rate of change in larval position from beginning of movement to end of track (χ2 Kruskal-Wallis (3) = 18.26, *p* = 0.00039, effect size (ε^2^) = 0.08, CI 95% [0.03, 1.00]). (D) Speed (scalar): total trajectory over time moved (χ2 Kruskal-Wallis (3) = 17.50, *p* = 0.000557, effect size (ε^2^) = 0.07, CI 95% [0.04, 1.00]). All parameters were calculated considering the position the larvae had at the moment of OMR initiation until reaching the end of the plate. Statistical analyses were performed using Kruskal-Wallis with Dunńs pairwise comparisons and Holm *p* adjustment method (Supplementary Table S13).

## Discussion

We developed a selectively CA-deficient zebrafish model by combining mutations in all genes involved in L-DOPA synthesis, *th, th2* and *tyr*. Surprisingly, our data indicate that Tyr activity contributes to more than half of L-DOPA to total DA synthesis in larvae. TH activity also contributes nearly half of total DA, while TH2 contributes only between 5% (compared to WT) and 15% (in *th* mutants). The strong contribution of *th* correlates with the broader expression of *th* compared to *th2*(Chen et al., 2009; Filippi et al., 2010; Yamamoto et al., 2010), and with higher total *th* mRNA expression(Chen et al., 2009). Previous *th* knock-down experiments reported smaller contributions of *th*(Formella et al., 2012), which may have been caused by partial Morpholino knockdown. Our results also demonstrate that TH2 indeed has Tyrosine hydroxylase(Chen et al., 2016; Yao et al., 2016), but not Tryptophan hydroxylase activity as previously suggested(Ren et al., 2013). Surprisingly, Tyr activity contributes to about one third of total DA. The relatively strong contribution of Tyr activity in zebrafish larvae is in contrast to data from mice, which showed that in TH-null mice only 2-3% of WT DA levels are detected in the brain, and analysis of TH-null TYR-null albino mice revealed that 80% of this residual DA originates from TYR activity(Rios et al., 1999).

Our analyses so far reveal no major anatomical differences between WT and *th, th2, tyr* triple mutant nervous system development. Importantly, it appears that the full complement of catecholaminergic neurons develops and project axons. CA depletion also does not appear to affect global brain axonal scaffold formation, and anatomically normal DA projections develop, including the long diencephalospinal DA tracts and peripheral projections to sense organs. Similarly, an anatomically normal midbrain dopaminergic system has been reported for dopamine deficient mice(Zhou et al., 1995). Synaptic activity plays a pivotal role in neuronal survival by coupling neurotransmitter release with neurotrophin uptake(Brady and Morfini, 2010). We infer that, despite lacking CAs, release of second transmitters such as glutamate or GABA(Filippi et al., 2014) may support survival and synapses maintenance in zebrafish CA neurons. Thus, our model enables future analyses of the impact of lack of CA neuromodulation on specific neural circuits, but, importantly, also new insights into non-CA activities of CA neurons, given that selective CA depletion may not eliminate other functions mediated by GABA or glutamate second transmitter systems in these CA neurons.

### Neural development in CA deficient larvae

DA has also been previously suggested to be required for proper development of zebrafish primary and secondary spinal motor neurons(Reimer et al., 2013). However, we find no differences in spinal motor neuron numbers in CA-deficient larvae, indicating that DA is not necessary for motor neuron development at larval stages. The postulated effects of DA on motoneuron development were suggested to be triggered by DA binding to D4a, a D2-like receptor, reducing cAMP levels and PKA activity, a negative regulator of hedgehog signaling, and thus activating hedgehog signaling that stimulates motor neuron development(Hammerschmidt et al., 1996; Lewis and Eisen, 2001). Considering that cAMP and PKA are regulated by several hormones and neurotransmitters(Sassone-Corsi, 2012), we propose that functional redundancy of GPCR and metabolic signaling networks may compensate for loss of DA in our genetic model, resulting in normal motor neuron development.

CAs have also previously been implied in mammalian stem cell systems, including sympathetic nervous system CA signaling in hematopoietic stem cell specification(Fitch et al., 2012). However, previous work already showed that in zebrafish earlier neural crest derived signals rather than sympathetic CA may induce hematopoietic stem cell specification(Damm and Clements, 2017), consistent with our finding of apparently normal blood cell development (data not shown). Therefore, while CAs may contribute to defined cell specification events, in absence of CAs this requirement may be compensated by parallel signaling pathway activities.

### CAs in control of hatching

Another physiological parameter affected by CA-depletion is HGC release. According to Schoots et al. (1983), dopamine indirectly inhibits hatching enzyme secretion by controlling prolactin release through the binding to D2-like receptors on lactotrophic cells(Schoots et al., 1982; Schoots et al., 1983). However, our finding of HGC retention in CA-deficient larvae indicates additional levels of CA-dependent direct HGC regulation. Even though HGCs do not express D2-like family receptors, they do express high levels of *drd5a* and *adrb3a*, as well as lower levels of *adrb1* and *adrb2a* (Farrell et al., 2018). Both, D1-like family receptors and β-adrenergic receptors are stimulatory G protein-coupled receptors (GPCRs), which activate the cAMP/PKA signaling cascade regulating secretion(Lopez and Gomez, 2010). Thus, our results suggest the presence of a parallel pathway with direct and positive control of CAs on HGCs, using potentially both DA and NA. A direct action of CAs on HGC secretion is also supported by our observation that *tyr* single mutants show enhanced HGC retention, with Tyr more likely acting through CA tissue levels rather than the hypothalamic-pituitary axis. We note that with respect to temporal control of development, melanocyte differentiation and Tyr expression correlate with HGC release in larval development. Thus, HGC release may not, or not exclusively, be regulated by neuronal CAs.

### Cardiovascular function in CAs deficient larvae

Similar to mammals, zebrafish cardiovascular function is regulated by CAs and highly sensitive to adrenergic drugs(De Luca et al., 2014). Since zebrafish are ectothermal, metabolic and physiological processes depend directly on environmental temperature(Seebacher, 2009). HR increases in direct relation with temperature(Gierten et al., 2020), which we also observed for all mutant genotypes. Despite zebrafish being ectothermal, their sinoatrial nodes have structurally and molecularly conserved features compared to mammals(Tessadori et al., 2012), whose heartbeat is generated by autorhythmic cardiomyocytes of the sinoatrial node, which do not depend on CAs. When we analyzed HR in individual CA-depleted genotypes across all temperatures, we observed a small but significant HR reduction, which at a specific temperature appears significant only at 36°C. This may reflect a requirement for activation of sympathetic NAergic components, as the effect we observe is similar in strength to HR reductions reported for *adrb1* adrenergic beta receptor mutants(Joyce et al., 2022).

Regarding to HRV, we observed that compared to standard temperature (28°C) both temperature decrease and increase cause an increment in HRV (Supplementary Figure S3B). Surprisingly, *th* mutants at low temperature show lower HRV and thus higher rhythmicity compared to *tyr* control siblings. Reduced HRV in humans is usually linked to cardiovascular disease and is used as a clinical marker to assess autonomic dysregulations. A dynamic balance of vagal and sympathetic input controls HRV, predominantly modulated by cholinergic and noradrenergic activities(Camm et al., 1996). Given that the noradrenergic component is depleted in our model, our results potentially reflect the diminished adaptability of CA mutants to changes in environmental conditions.

### Motor behavior in CAs deficient larvae

DA is also an important modulator of central sensory and motor systems. As in mammals, zebrafish DA systems modulate behavioral responses, including locomotor development(Lambert et al., 2012) and sensory processing(Mu et al., 2012). In concordance with these findings, our results showed that loss of TH activity elicits a reduction in spontaneous swimming. The basic commands for executing rhythmic locomotor outputs rely on spinal central pattern generator activity(Wiggin et al., 2012), which is subject to refinement by supraspinal modulatory inputs, including DA (Hachoumi and Sillar, 2020). In zebrafish, the diencephalospinal DA tracts(McLean and Fetcho, 2004; Tay et al., 2011) project exclusively from posterior tubercular DC2 and DC4 DA neurons, which solely express *th* but not *th2*, and correspond to the mammalian A11 DA system. Our results are concordant with experiments carried out on 5 dpf larvae, where laser ablation of DC2 and DC4/5 groups elicited a reduction in swimming behavior(Jay et al., 2015).

Interestingly, in larvae lacking TH2 activity, we did not observe an effect on larval locomotion. In contrast, McPherson et al.(McPherson et al., 2016) reported that chemogenetic ablation and optogenetic activation of *th2* neurons at 8 dpf affect swim bout initiation frequency. This apparent incongruence with our results could be explained by overlapping expression of *th* and *th2* in the DC7 DA group and in *th2* neurons of the posterior recess(Filippi et al., 2010). Dopamine transporter may mediate uptake of DC7 *th* derived DA into posterior recess *th2* neurons in *th2* mutants, potentially rescuing some activities of *th2* DA neurons. Therefore, our experiments may not allow to define *th* or *th2* specific phenotypes, but rather reflect phenotypes caused by decrease of total interstitial DA levels in the brain. Further, chemogenetic ablation or optogenetic activation of posterior recess *th2* neurons(McPherson et al., 2016) abolishes both DA and second transmitter, likely GABAergic, activity of these neurons, while in *th2* mutant embryos we expect second transmitter systems in these neurons to still be active.

Upon visual stimulation with moving gratings, *th* mutants showed an abnormal OMR, including reduced directionality, speed, and velocity, while latency in response was considerably increased. Since the execution of optic-flow oriented movement requires the engagement of both sensory and motor pathways, lack of DA signaling may differentially affect the kinematic outputs of OMR. Among the four parameters tested, directionality and latency rely on proper visual information processing in the retina and pretectum, while latency, speed and velocity are also encoded in motor systems. In mammals, both bright and dim light have been shown to induce DA release by DA-amacrine cells (DA-AC), which improve visual and contrast sensitivity(Dowling, 1991; Jackson et al., 2012). The zebrafish retina contains DA-AC as well as D1- and D2-like receptor expressing cells(Boehmler et al., 2004; Li et al., 2007), and DA-AC ablation induces impairments in visual sensitivity, effects that were reversed by DA-agonists treatment(Li and Dowling, 2000). Thus, we hypothesize that lack of DA-AC *th* expression will impair the proper activity of retinal cell types, leading to increased visual thresholds. Retinal Ganglion Cell (RGC) firing may only be produced after consecutive stimulation, delaying the response to stimulus onset, contributing to the increased latency in *th* mutants. Likewise, lack of DA modulation of direction selective RGCs could also affect the extraction and transfer of motion direction to pretectal arborization fields(Kramer et al., 2019), resulting in a reduced directionality. Response latency may also involve DA modulation of supraspinal circuits controlling motor initiation. On the other hand, we suggest that the reduction in velocity and speed observed in *th* mutants may be related to dysfunction in downstream motor control circuits, which compromise the execution of the response. This is concordant with previous D1 agonist treatments performed in zebrafish larvae, which elicited faster swim responses upon OMR, based on recruitment of more primary and secondary motor neurons controlling fast versus slow swim responses(Jha and Thirumalai, 2020).

## Conclusion

We think that our CA-deficient zebrafish model will be instrumental to address unresolved issues on DA systems development and circuit maturation, and help to distinguish between DA/NA specific effects and other second transmitter activities of CA systems. We note that the rather subtle phenotype of the triple mutant larvae was surprising to us. We validated based on published ssRNAseq data(Raj et al., 2020) that DAergic and adrenergic receptors are already broadly expressed in most brain regions at 5 dpf stage (Supplementary Figure S6). In accordance, effects of pharmacological manipulation of CA systems have been reported in 5 dpf or earlier larvae(Jay et al., 2015; Joyce et al., 2022; Randlett et al., 2019; Schoots et al., 1983; Thirumalai and Cline, 2008). Thus, a lack of more severe phenotypes is not caused by immature CA systems. We speculate that circuit development in the absence of CAs may lead to more pronounced compensation of loss of DA by other neuromodulators like serotonin. Serotonin has also been shown to pattern locomotor network activity in zebrafish(Brustein et al., 2003). In mammalian systems, DA and serotonin have also been shown to synergize or partially compensate in control of other behaviors including reward(Fischer and Ullsperger, 2017). The complex interactions of DA, serotonin and other neuromodulators like endocannabinoids and other neuropeptides(Peters et al., 2021) may explain how potential differences in circuit development as well as intrinsic compensation mechanisms in neuromodulation may cause rather normal behavior in the absence of DA. Our model may thus also provide new avenues of research on the interplay of neuromodulators in normal and diseased states, including Parkinson’s and depression.

## Materials and Methods

### Zebrafish maintenance, breeding and transgenic lines

Zebrafish (*Danio rerio*) embryos and larvae were obtained by natural breeding and staged accordingly(Kimmel et al., 1995). Embryos were incubated at 28.5°C in E3 (5 mM NaCl, 0.17 mM KCl, 0.33 mM CaCl2, 0.33 mM MgSO4, pH 7.0) with 0.01% methylene blue medium at a 14:10 light/dark photoperiod. Wild type zebrafish are from an AB/TL genetic background. The generation of the zebrafish lines *th^m1403^*and *th2^m1420^* is described in this paper. Tyrosinase mutants (*tyr^tk20^*) were kindly provided by Herwig Baier(Page-McCaw et al., 2004). Triple mutant embryos were obtained by incross of heterozygous genotypes for *th^m1403^* and *th2^m1420^*, and homozygous for *tyr^tk20^*. The following transgenic lines were used: *Tg(th:Gal4-VP16)^m1233^*, *Tg(UAS:eGFP-CAAX)^m1230^*(Fernandes et al., 2012) and *Tg(mnx1:GFP)^ml2Tg^*(Flanagan-Steet et al., 2005). All zebrafish care and experiments were conducted according to the German Animal Welfare Laws.

Nomenclature: We note that by zebrafish nomenclature conventions (https://zfin.atlassian.net/wiki/spaces/general/pages/1818394635/ZFIN+Zebrafish+Nomenclature+Conventions), the triple mutant embryos, based on the chromosome locations of the loci (*th2* chromosome 4; *tyr* chromosome 15; *th* chromosome 25) should be written *th2^m1420/m4120^; tyr^tk20/tk20^; th^m1403/m1403^.* For easier reading, we use “*th, th2, tyr* mutants” or just “triple mutants”, and for double mutants e.g. “*th, tyr* (double) mutants”. In some figures we use *th th2 tyr* for mutant homozygous loci for sake of space. Heterozygous combinations are designated *th+/-, th2+/-, tyr+/-*.

### Generation of *th* and *th2* mutant alleles using TALEN

Loss of function mutations in the *th* and *th2* genes were induced using the TALEN technology(Huang et al., 2011). TALENs were designed using the online tool *“Mojo Hand”* http://www.talendesign.org(Bedell et al., 2012) and assembled using the Golden Gate cloning method(Cermak et al., 2011). The clones containing the full length TALENs were sent for sequencing and the results were analyzed using the online program TAL Plasmids Sequence Assembly Tool (http://bao.rice.edu/Research/BioinformaticTools/assembleTALSequences.html). TALEN plasmids were linearized and then transcribed using the mMESSAGE mMACHINE® High Yield Capped RNA Transcription Kit (Thermo Fisher). TALEN mRNA pairs were injected at 100 or 200 ng/μL each into one-cell stage wild type AB/TL embryos and developed for one day to perform DNA extraction and PCR amplification (Figures 1B, 2C). PCR products were then digested with TaqI or NsiI (Figure 2B,C) for *th2*, or restricted with EcoRI for *th* (Supplementary Figure 1B). Non-cleaved bands were sent for sequencing to validate TALEN efficiency.

For the *tyrosine hydroxylase* locus (*th*), the knock-out was carried out by disrupting an EcoRI restriction site at the beginning of the open reading frame (ORF) to facilitate the analysis through restriction fragment length polymorphism (RFLP). TALENS were designed to bind to 5’-TCCGCGCGCGCGCCATCTGA-3’ on the coding and to 5’-GCAGCTCCACATCTTCCACA-3’ on the complementary strand. The spacer region was *ac**atg**cc**gaattc**aa* carrying the restriction enzyme recognition site (*gaattc*) for EcoRI (Figure 1A). Disruption in the restriction site results in undigested bands that demonstrate a TALEN-induced in/del mutation (Figure 1B, EcoRI digestion, uncleaved). Sequencing of the intact bands revealed three different indel mutations. A 11 bp deletion (allele *m1404*) and a 1 bp insertion (allele *m1405*) each caused a frameshift and premature stop codon formation, while a 7 bp deletion (allele *m1403*) disrupted the native ATG initiation codon (Figure 1C). In this study the *th^+/m1403^* line was used.

The knock-out for *tyrosine hydroxylase 2* (*th2*) was obtained by complete deletion of the coding region, from exon 1 to exon 12 (Figure 2A). To achieve this goal, two pairs of TALEN targeting the first (*th2*-E1) and last (*th2*-E12) exon were designed. The first pair was directed towards the start of the ORF of the *th2* gene by targeting the sequences 5’-TAACTGGATATATTCA-3’ on the coding and 5’-CGGACAGTATAGCGC-3’ on the complementary strand of exon 1, while the second pair disrupted the end of exon 12 by binding to 5’-GAGGTCTAGGGTTGA-3’ and 5’-TCTCAGATGTAATAAAC-3’ on the coding and complementary strands respectively. TALENs binding regions include restriction sites for TaqI (exon 1) and NsiI (exon 12). TALEN activity was tested by RFLP assay. Digestion with TaqI (*th2*-E1) resulted in one thick band seen at ∼250 bp, which corresponds to the overlap of the two digestion products of similar size (242 bp and 250 bp, Figure 2B-C). Digestion with NsiI (*th2*-E12) breaks down the amplicon in two bands of 423 bp and 140 bp (Figure 2B,C). In TALEN-injected samples we observe both cleaved and uncleaved bands (*th2*-E1: 492 bp; *th2*-E12: 563 bp; Figure 2C), demonstrating the efficacy of TALENs in modifying the restriction sites. Combined activity of both TALENs results in a 10 kb deletion (Figure 2E), indicated by the presence of a 382 bp product (Figure 2D), which is only possible after the complete deletion of the E1 to E12 sequences (Figure 2B). In this study the *th2^+/m1420^* allele was used.

For each mutation, TALEN injected founders (G0) were raised and identified following outcross with AB/TL wild type fish. The resulting egg clutches were screened using PCR and restriction assay. For egg clutches with a mutant allele, siblings were raised as filial generation 1 (F1). F1 heterozygotes were crossed to AB/TL wild type fish to establish a stable mutant line, and the sequence for each allele determined (Figures 1C and 2E).

### Genotyping

Genotyping of larvae was done after completion of the experiments, using DNA isolated from either the whole larvae, or tail fin biopsies. Adult fish were anesthetized with MS-222 and genotyped from tail fin biopsies. After a 10 min incubation period at 98°C, samples were digested with Lysis Buffer (stock 10x: 100 mM Tris pH 8, 500 mM KCl, 3% Tween 20, 3% NP40, 10 mM EDTA in ddH_2_O) and Protein-kinase K (10 µg/ml in PBST, A3830, AppliChem) overnight followed by 10 min at 98°C. Mutants for *th^m1403^* were identified using primers flanking the 7bp deletion at the EcoRI site in the first exon (forward: 5’-AAGATGTTCAATGACGGAG-3’; reverse: 5’-GTTGAAGTGGACAATGTAG-3’), followed by a restriction assay with the enzyme EcoRI*-*HF (R3101, BioLabs) (Figure 1D). For *th2^m1420^* genotyping, a common reverse primer (5’-ATGCCCCTGGAAATAGGC-3’) and two mismatch forward primers hybridizing either the WT (5’-CTTGGGGAGACTGGGACAG-3’) or mutant (5’-CTGTCGCTTGAAGCACCTG-3’) sequences were used (Figure 2D), resulting in DNA fragments of 286 bp for WT and 396 bp for mutant alleles. Homozygous mutants for *tyr^tk20^* were identified by absence of melanocyte-derived dark pigmentation.

### Whole-mount *in situ* hybridization (WISH)

Digoxigenin-labelled antisense RNA probe for *th*(Holzschuh et al., 2001) and *hgg1* (*hatching gland 1* or *ctslb cathepsin Lb*)(Thisse et al., 1994) were *in vitro* synthetized using the mMESSAGE mMACHINE T3 Transcription Kit (Ambion) and detected using Anti-digoxygenin-alkaline-phosphatase (0,2 µl/ml PBST, #11093279910, Roche). Whole-mount *in situ* hybridization (WISH) was performed at 4 days post fertilization (dpf) as described(Thisse and Thisse, 2008).

### Immunofluorescence Imaging

Immunofluorescence staining for GFP detection was carried out using chicken anti-GFP (1:400 IgY, A10262, Life Technologies) and Alexa Fluor 488 goat anti-chicken (1:500, Fab fragment, IgG, A11039, Life Technologies). Spinal cord projecting neurons (Mauthner cells) were detected using mouse anti-neurofilament 3A10 IgG (1:20, supernatant, DSHB, 3A10)(Dodd and Jessell, 1988) with Alexa Fluor 555 goat anti-mouse as secondary antibody (1:500, IgG, A21425, Life Technologies). Samples were counterstained with nuclear stain TOTO-3 iodide (T3604, Invitrogen) for general anatomical orientation. Anti-TH immunofluorescence was carried out with a polyclonal Rabbit anti zebrafish TH antibody(Ryu et al., 2007).

### Imaging

WISH stained larvae were mounted in 100% glycerol and imaged with an Axioplan Microscope (10x/NA0.3 objective, Carl Zeiss, Germany) with AxioVision Software (Carl Zeiss, Germany), typically recording Z-stacks of the brain. Z-stacks were processed in ImageJ (Fiji Edition, https://imagej.net/software/fiji/) to generate minimum intensity projections. Whole mount immunofluorescence of *Tg(mnx1a:GFP)* labelling spinal motor neurons were imaged in 80% glycerol with an Examiner microscope (20x/NA1.0 water immersion objective, Carl Zeiss, Germany) using the ZEN blue software (Carl Zeiss, Germany). Z-stacks of individual somites were analyzed with the 3D function. Motor neurons were counted at 33 hours post fertilization (hpf) in 4 somites (7, 8, 9 and 10) of 5 embryos each, while at 3 dpf 3 somites (7, 8 and 9) were counted in 5 embryos each. Immunofluorescence performed on CA-tracts and axonal projections were mounted in 1% agarose containing Vectashield Mounting Medium (H-1000, Vector Laboratories), and imaged using an inverse LSM880 laser scanning confocal microscope (25x/0.8NA multi-immersion objective, Carl Zeiss, Germany).

### Dopamine ELISA

For dopamine quantification, 5 dpf larvae were euthanized with MS-222. Two sampling procedures were used: (1) heads of WT AB/TL and *tyr-/-* larvae at 20 hpf or 5 dpf were collected and immediately pooled before freezing (designated “heads frozen in bulk”) in 500 µl of Homogenizing Medium (HM, 0.01N HCl; 1 mM EDTA; 4 mM Sodium Metabisulfite; in 1x PBS), and stored at -80°C. (2) For 5 dpf progeny obtained from single (for *th* or *th2*), double (for *th th2*) or triple (for *th, th2, tyr;* parents homozygous for *tyr*) heterozygous parents, heads were collected and stored separately (“single heads”). Tails were cut off for genotyping, while heads were transferred to 50 μl of HM and stored at -80°C.

The observed fraction of specific genotypes in these crosses correspond to the expected genetic Mendelian segregation. 50 homozygous triple mutants were obtained from 1187 pre-sorted *tyr-/-* larvae genotyped for *th* and *th2* (4.7 % double homozygous, expected 1/16 = 6.25%). From *th+/-th2 +/-* crosses, 1200 larvae were genotyped for *th* and *th2* with 4,3 % double homozygous detected (expected 6.25%). From *th+/-* crosses, out of 480 larvae analyzed 24.8 % *th-/-* mutants were recovered (expected 25%), and from *th2+/-* crosses out of 415 larvae 25.5 % *th2-/-* mutants were recovered (expected 25%).

After obtaining the genotyping results from tail tissue, 50 heads of the specific desired genotype were pooled to a final volume of 500 μl. Samples from (1) and (2) each were manually homogenized on ice using a micro pestle in a 1.5 ml reaction vial. Samples were then placed in boiling water for 3-4 minutes and centrifuged for 5 minutes at 20.000 rpm, 4°C. Dopamine-containing supernatants were collected and stored at -80°C. Extraction efficiency was compared using Bradford Assay (Protein Assay, #5000006, Bio-Rad®). As expected, WT 20 hpf stage extracts had lower protein amounts given that the cell number in embryos at 20 hpf is much smaller than at 5 dpf. Dopamine quantification was carried out using a competitive Enzyme-Linked ImmunoSorbent Assay (Dopamine Research ELISA™ Kit, BA E-5300, LDN, Nordhorn, Germany). Extraction and acylation of dopamine were performed in a sample volume of 500 μl, followed by enzymatic conversion and dopamine ELISA. All the steps involved in dopamine quantification were done following the manufactures’ specifications. The ELISA was validated by 1:2 dilutions of WT 5 dpf, WT 20 hpf and *tyr* mutant samples as well as by 1:4 dilution of WT 5 dpf. For the 20 hpf WT sample, 150 whole zebrafish larvae were used instead of 50 heads, to get readings closer to the linear range of the ELISA; the calculated DA amounts were divided by three to obtain the data on DA content in 50 larvae. In addition, the ELISA was validated by spiking 1 ng or 10 ng DA (Dopamine HCl Sigma H8502) into WT 5 dpf samples obtained from 50 WT bulk heads (For data see Supplementary Table 1). The ELISA readings indicate that under our extract conditions, 1 ng DA caused higher and 10 ng DA lower readings than expected (Supplementary Table 1). Therefore, the ng DA per 50 heads values in Figure 3 may be lower than actual.

Comparisons were performed among samples obtained with the same extraction method. The bulked head samples are expected to better represent the total DA in zebrafish larval heads, because in single head preparation DA may leak out into freezing media that later was discarded to limit total sample volume to 500 µl. Indeed, comparing WT 5 dpf samples, DA amounts measured in single head extracts were smaller than those from bulked heads. Given that protein extraction efficiency in *th th2* double and *th th2 tyr* triple mutants was similar to WT single head samples, we think that the extraction method does not significantly affect DA measurements in these samples compared to WT.

### Heart rate

Heart rate (HR) and heart rate variability (HRV) were obtained as described(Gaur et al., 2018; Shaffer and Ginsberg, 2017). Briefly, heartbeat was recorded at 3 dpf between 76 and 81 hrs. post fertilization in non-anaesthetized and free moving larvae. Video acquisitions were done at 35 frames per second (fps) using a USB 3.1 monochrome camera (DMK 37BUX250, Imaging Source, Germany) equipped with the IC Capture 2.4 software. Recordings were performed in larvae at control temperature (28°C) and at four thermal challenges (20°C, 24°C, 32°C and 36°C). Larvae were all raised at 28°C, transferred to the required experimental temperature, given a 1 min period of acclimatization, and subsequently heart beat was recorded on video of 1 min. Video files were saved in AVI (Audio Video Interleave) format and decompressed using FFmpeg(Tomar, 2006). All spontaneous motion such as pectoral fin and gill movements were not considered for the final analysis. Video frames were imported into ImageJ. Heart rate was extracted with ImageJ by selecting a region in the heart(Gaur et al., 2018) and using the “plot Z-axis profile” function in ImageJ to reveal the pixel intensity changes over time produced by every heartbeat, while HRV was calculated as the root mean square of the successive differences between heart beat intervals (RMSSD)(Shaffer and Ginsberg, 2017).

### Hatching phenotype

Hatching gland cell (HGC) retention was assessed in larvae from crosses of *th+/-, th2+/-, tyr-/-* parents. At 3 dpf the larvae were divided in two groups based on the presence (HGC+) or absence (HGC-) of HGCs in the pericardial membrane (Figure 6A). About 25% of the larvae still had HGCs at 3 dpf (HGC+ phenotype) and were selected, while a similar number of HGC-larvae were randomly chosen from the remaining three quarters of larvae. After genotyping, the relation between HGC-phenotype and genotype was examined. At 5 dpf the abundance of granule containing cells in the pericardial membrane was classified into high (over 80% of pericardial membrane still containing HGC), low (less than 20% of the pericardial membrane contain HGC) and no-retention (no HGC cells observed) (Figure 6D). The HGC presence was assessed using a binocular stereomicroscope (Leica MZ12, Germany).

### Transcriptomic Analyses

scRNAseq data from 5 dpf zebrafish brain and eye cells(Raj et al., 2020) were downloaded from GEO: GSE158142 (https://www.ncbi.nlm.nih.gov/geo/query/acc.cgi?acc=GSE158142, GSE158142_zf5dpf_cc_filt.cluster.rds, downloaded on 05/11/2020) and analyzed by the Seurat package (v4.3.0)(Satija et al., 2015). Seurat object was updated from v2.X to v3.X using the SeuratObject::UpdateSeuratObject (v4.1.3) function. Clustering and Annotation was retained from original data from Raj et al., 2020. The average expression levels for the respective genes were calculated using Seurat::AddModuleScore. FItSNE plots were generated using Seurat::DimPlot and Seurat::FeaturePlot functions. The expression of CA Receptors in HGCs was investigated using the online resource Daniocell (https://daniocell.nichd.nih.gov/index.html, v1.1, 02/01/2024)(Sur et al., 2023). HGC clusters (axia_15, axia_18, axia_20) were confirmed by expression of *cathepsin Lb (ctslb)* and *hatching enzyme 1* (*he1.1)*.

### Spontaneous Locomotor Activity

Video recordings of spontaneous swimming were performed under white light at a frame rate of 30 fps for 1 hour in the Noldus DanioVision observation chamber (model number DVOC004x/T; Noldus, Wageningen, Netherlands) at 28.5 °C in a 24-well format. Visual recognition of unpigmented larvae in the videos was improved by measuring locomotor activity on a 24 multi-well plate lid (lid with circular lines for each well; Cellstar, Grainer Bio-One, Germany), instead of the deep well bottom parts of the plastic dishes. The lines on the lid separating the wells were made into circular barriers by application of 3% agarose outside the circles to avoid mixing of larvae. The lid wells had 12 mm diameter and 7 mm depth and were each filled with 800 μl E3 medium. Videos were recorded using Noldus DanioVision standard setting and tracked using the software Ethovision XT14.

### Optomotor Response (OMR)

OMR assays were performed in a custom-made multi-chamber plate containing 6 rectangular rails of 1 cm width and 7 cm length (Figure 8B). The transparent plate was positioned directly on a 4K 15,6 Zoll Portable Monitor (3840 * 2160 pixels; Thinlerain Shenzhen Xinliangyuan Technology Co., Ltd. Limited, Shenzhen CN). The black and white OMR gratings were generated with the Visual Stimulator made by Prof. Michael Bach (https://michaelbach.de/sci/stim/movingGrating/index.html;(Rainy et al., 2016)) using maximum contrast. Bright light for pre-stimulus conditioning was followed by presentation of moving gratings from below to free-swimming larvae. For assessing OMR, *tyr* phenotype zebrafish larvae were acclimatized to bright light during 30 min in order to avoid visual impairments related with Sdy/Tyrosinase depletion(Page-McCaw et al., 2004). Movement tracks were manually generated from the video data using the Manual Tracking plugging in ImageJ. The animals were classified as responders (engage OMR), non-responders (move randomly), or non-moving (do not move at all). For each responder, the OMR was individually measured and trajectory, displacement and time were extracted from the tracks. The measurement started with the onset of the stimulus and ended when the larva reached the end of the trail. In order to make the results comparable, raw variables were normalized considering the ratios among them, resulting in directionality (displacement/trajectory), speed (trajectory/time) and velocity (displacement/time). Latency was measured as the time delay between stimulus onset and optic-flow oriented movement.

### Statistics

Statistical analysis and plots were done using the R package ggstatsplot(Patil, 2021). Data distribution was tested using Shapiro-Wilk test for small sample size (n < 20) and Kolmogorov-Smirnov normality test for larger sample sizes (“ks.test”). Parametric and non-parametric statistical tests were selected according to data distribution, followed by post-hoc pairwise comparisons. The function “ggbetweenstats” was used for statistical testing of unpaired samples as well as for effect size estimations. For categorical data, the function “ggbarstats” was used. Differences among qualitative variables were assessed using Pearson’s chi-squared (χ^2^) test. Two-way ANOVA was performed to evaluate the effect of temperature and genotype on heart function. Since ggstatsplot does not support two-way ANOVA, the test was performed using the functions “aov” and “Anova” with Type-III sums of squares for unbalanced designs (different samples size within each level). “TukeyHSD” was performed post-hoc for multiple pairwise comparisons. Statistical significance was set at p < 0.005. Data presented as violin plots depict data distribution from maximum to minimum values as well as boxplots showing median, upper and lower quartiles, while bar plots show mean and standard deviation (SD). For outlier analysis, data values differing significantly from the observations were identified according to Tukey’s rule using an outlier coefficient of 1.5. To reduce the probability of type II error, in Supplementary Figure S5 outliers were eliminated from the analysis. All experiments and analysis were performed blinded to the genotype. Further statistical details including tests and sample sizes are presented in the figure legends.

## Supporting information

PDF with assembles supplementary figures

## Data availability

All numerical analyses of data are provided in the supplementary tables. All primary image data, videos and behavioural tracking are available on request.

## Acknowledgements

We thank Sabine Götter for excellent fish care. Funded by Baden-Württemberg Landesgraduiertenförderung doctoral stipend to SPZ, the Deutsche Forschungsgemeinschaft (DFG, German Research Foundation) under Germany’s Excellence Strategy—EXC-2189— Project ID: 390939984, and by DFG—322977937/GRK2344 MeInBio.

## Author contributions

S.P.-Z. designed and performed experiments, analyzed data, and wrote and assembled the manuscript; R.P. designed and performed experiments, analyzed data, assembled figures, and wrote parts of the manuscript; K. Ø. and G.A. performed genetic experiments; D.F. and J.O. designed and performed experiments, analyzed data; J.H. T.S. and F.V. designed and supervised experiments and contributed to data analysis; W.D. designed the study, supervised experiments, analyzed data, obtained funding, wrote parts of and edited the manuscript.

