## Supplementary material for "A genetic model for development, physiology and behavior of zebrafish larvae devoid of catecholamines": PDF with assembles supplementary figures

**Supplementary Data**

**Supplementary Figures (Pages 2 – 11)**

Supplementary Figure S1. Catecholamine biogenesis and catecholamine systems in zebrafish (Page 2)

Supplementary Figure S2. Expression of HGC markers and selected CA Receptors in HGC clusters (Page 4)

Supplementary Figure S3. Comparison of HR and HRV changes at different temperatures for specific *tyr, th* and *th2* mutant combinations (Page 5)

Supplementary Figure S4. Spontaneous locomotion of *AB/TL* WT controls and *tyr* mutant larvae (Page 6)

Supplementary Figure S5. OMR kinematic parameters in CA depleted larvae (with outlier analysis) (Page 7)

Supplementary Figure S6. Expression of CA receptors in 5dpf brain cells (Page 8)

**Supplementary Tables (Table Legends Pages 10-11)**

(One Supplementary EXCEL File with 13 Sheets for Tables S1 - S13)

Table S1: Figure 3 Dopamine ELISA

Table S2: Figure 4C *th* cell counts

Table S3: Figure 5B cell counts 33 hpf

Table S4: Figure 5B cell counts 3 dpf

Table S5: Figure 6B HGC phenotypes 3 dpf

Table S6: Figure 6E levels of HGC retention among genotypes from 3 dpf to 5 dpf

Table S7: Figure 7HR and HRV temperature dependence data

Table S8: Figure 7 HR and HRV temperature dependence statistical analysis

Table S9: Figure 8A locomotor activity

Table S10: Figure 8A locomotor activity

Table S11: Figure 8C optomotor response

Table S12: Figure 9 optomotor response

Table S13: Figure 9 optomotor response two-sample Kolmogorov-Smirnov test

**Supplementary References (Page 12)**

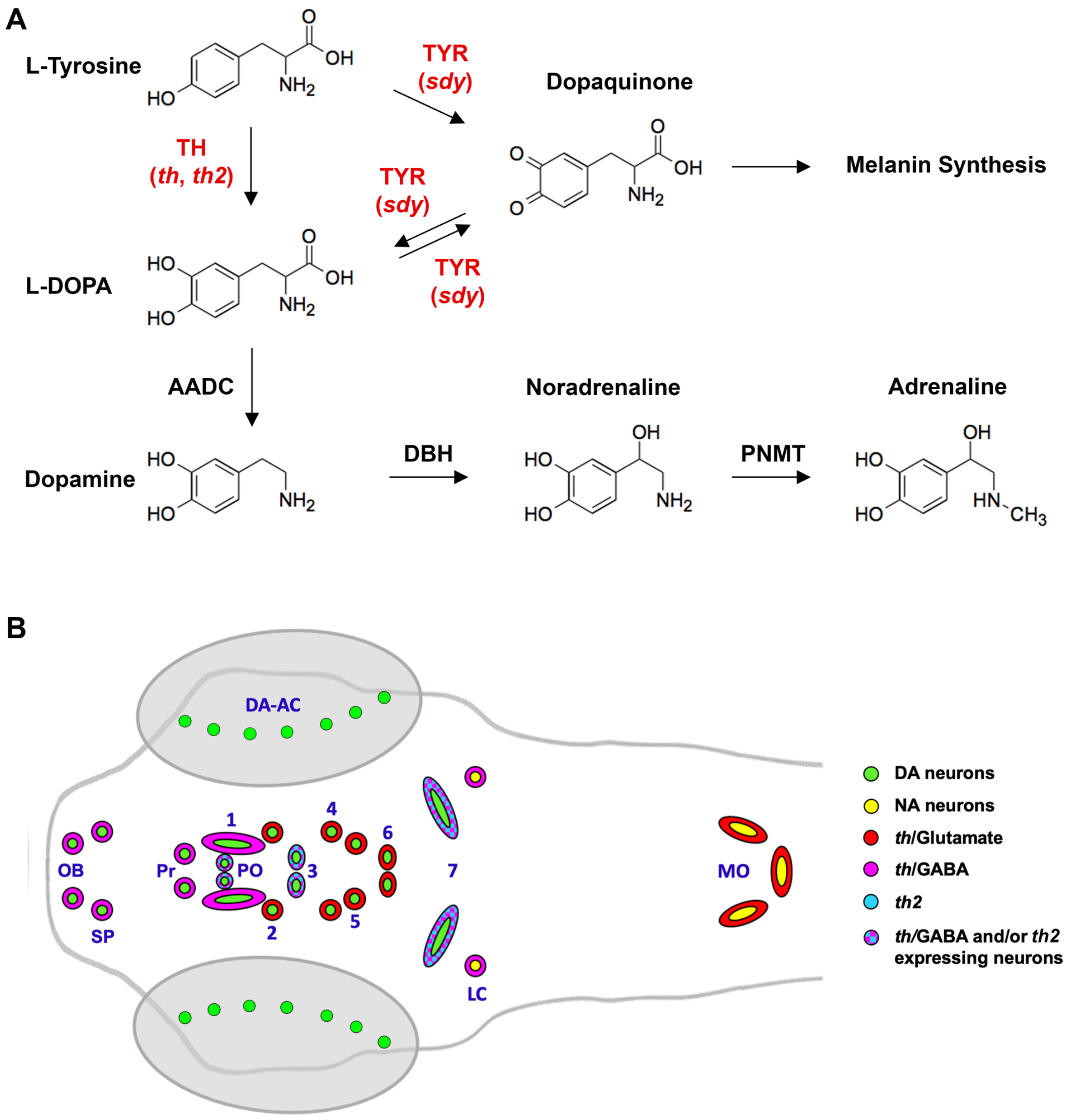


**Supplementary Figure S1. Catecholamine biogenesis and catecholamine systems in zebrafish**

(A) Enzymatic pathways of catecholamine synthesis. AADC: Aromatic L-Amino Acid Decarboxylase; DBH: Dopamine β-hydrolase; L-DOPA: L-3,4-dihydroxy-phenyalanine; PNMT: Phenylethanolamine-N-methyltransferase; TH: Tyrosine hydroxylase (*th* and *th2* loci in zebrafish); TYR: Tyrosinase (*sdy* locus in zebrafish). Modified from(Eisenhofer et al., 2003). (B) Catecholamine systems in zebrafish larvae. Anatomical scheme of CA systems in the 3-5 days post fertilization (dpf) larval zebrafish brain. Dorsal view, anterior left. CA systems expressing *th* are shown using the nomenclature developed by Rink et al.(Rink & Wullimann, 2001; Rink & Wullimann, 2002). The DA systems have evolved significantly and show both distinct and shared features with the mammalian systems(Smeets & Reiner, 1994; Yamamoto & Vernier, 2011). Compared to mammalian DA systems, the diencephalospinal DA systems (A11 in mammals, DC 2,4,5 and 6 in the zebrafish posterior tuberculum; DC =

DA Cluster, figure shows numbers only), the prethalamic system (A13 versus DC1), olfactory system (A16 versus OB), and amacrine retinal system (A17) are considered largely conserved. In zebrafish, the preoptic system (A15 versus PO) is complemented by the optic recess region DA system (ORR). The hypothalamic A12 and A14 systems develop divergently in the zebrafish, which have mammillary (DC3) and tuberal (DC7) hypothalamus DA systems(Schredelseker et al., 2020). Zebrafish, contrary to mammals, have DA systems in the pretectum (Pr) (Holzschuh et al., 2001) and extended medial amygdala of the subpallium (SP) (Armbruster et al., 2025). Furthermore, mesdiencephalic DA clusters (A8, A9 and A10 in mammals) are absent in zebrafish, likely due to secondary loss in teleost evolution, given that they are found in lungfish(Reiner & Northcutt, 1987), and very few non-functional remnants have also been detected in zebrafish(Altbürger et al., 2023). The hindbrain (LC locus coeruleus and MO medulla oblongata) and peripheral (carotid body and sympathetic ganglia; not shown) NA systems are largely conserved(Ma, 1994; Ma, 1997; Holzschuh et al., 2001).

The CA clusters DA (green) and NA (yellow) have initially been described based on *th* expression. By 5 dpf, *th2* (cyan) expression domains occur in the hypothalamic posterior and lateral recess, the paraventricular organ and preoptic region(Filippi et al., 2010; Yamamoto et al., 2010). GABAergic (magenta) or glutamatergic (red) second transmitters are indicated. Abbreviations: AC: amacrine cells of retina; OB: olfactory bulb; SP: subpallium; Pr: pretectum; PO: Preoptic system; 1-7: DC clusters 1 to 7; LC: locus coeruleus; MO: medulla oblongata.

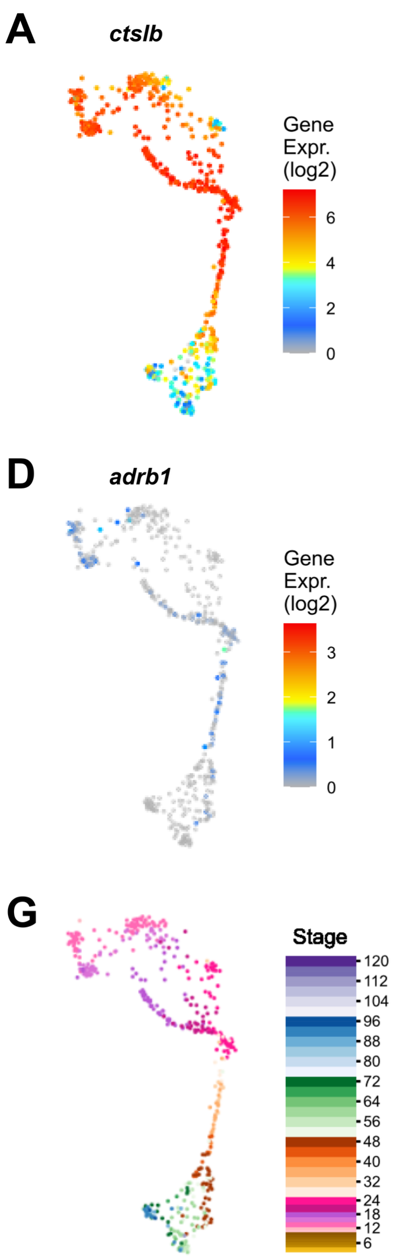


**Supplementary Figure S2. Expression of HGC markers and selected CA Receptors in HGC clusters**

Expression of HGC markers (A,B) and selected CA Receptors (C-F) in HGC clusters of scRNAseq data from Daniocell online resource (https://daniocell.nichd.nih.gov/index.html, v1.1). Data are from 3.3–120 hpf (G) whole-animal AB/TL zebrafish. Axial mesoderm clusters axia_15, axia_18, and axia_20 are shown.

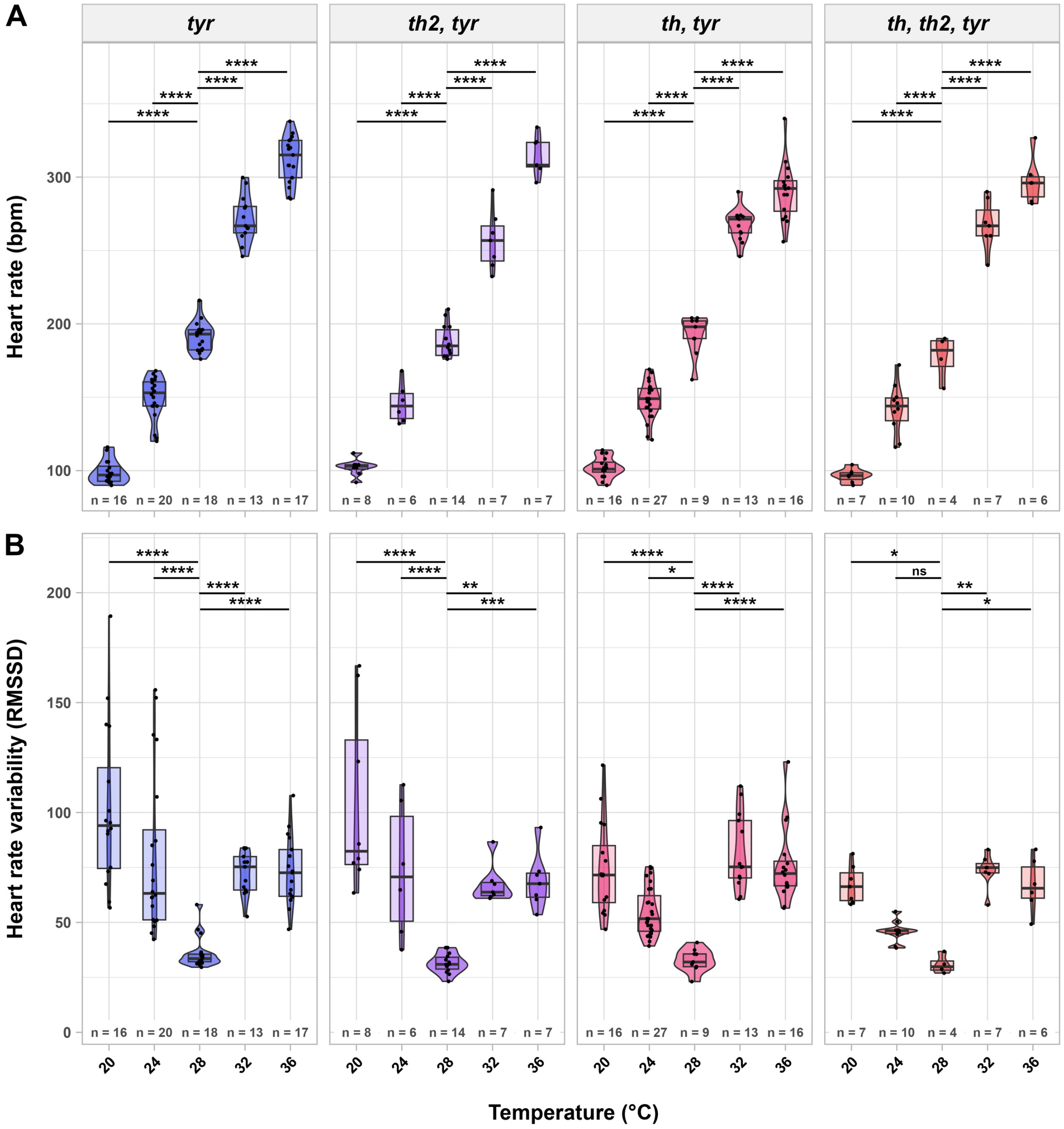


**Supplementary Figure S3. Comparison of HR and HRV changes at different temperatures for specific *tyr, th* and *th2* mutant combinations**

Relates to Main Figure 7. (A) HR and temperature appear positively related for all genotypes. (B) HRV increases with temperature challenges. Significances were calculated comparing each temperature with 28°C. Statistical analysis was performed using two-way ANOVA with Tukey's multiple pairwise-comparisons test (See supplementary table S8). *: p-value < 0.05; **: p-value < 0.001; ***: p-value < 0.0001; ****: p-value < 0,00001.

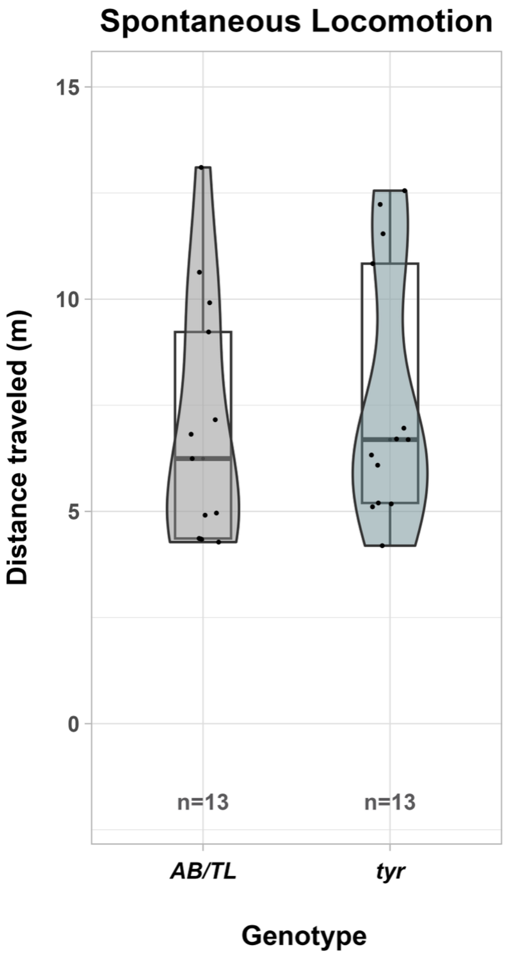


**Supplementary Figure S4. Spontaneous locomotion of *AB/TL* WT controls and *tyr* mutant larvae**

Relates to Main Figure 8. Spontaneous locomotor activity measured as distance traveled in 1 hour (hr) is not significantly different in *tyr* mutants compared to AB/TL control larvae. Statistical analysis was performed using two-tailed Mann-Whitney test (W = 68, $p$ = 0.41, effect size (r (rank biserial)) = -0.20, CI 95% [-0.57, 0.25])

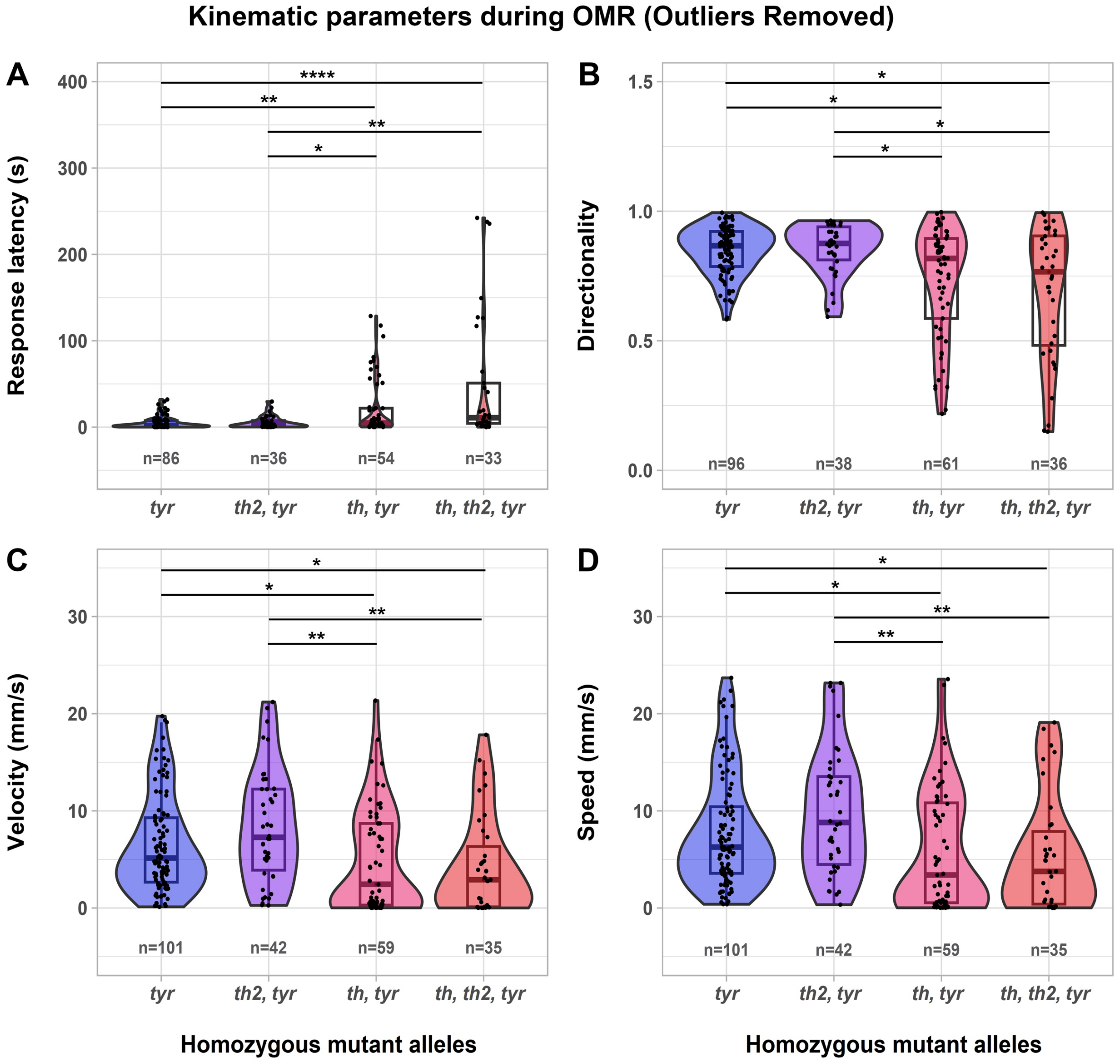


**Supplementary Figure S5. OMR kinematic parameters in CA depleted larvae (with outlier analysis)**

(A) Latency in OMR initiation ($\chi 2$ Kruskal-Wallis (3) = 25.33, $p$ = 0.00001, effect size ($\varepsilon^2$) = 0.12, CI 95% [0.05, 1.00]), (B) Directionality ($\chi 2$ Kruskal-Wallis (3) = 15.78, $p$ = 0.00126, effect size ($\varepsilon^2$) = 0.07, CI 95% [0.04, 1.00]), (C) Velocity (vector) ($\chi 2$ Kruskal-Wallis (3) = 21.13, $p$ = 0.00009, effect size ($\varepsilon^2$) = 0.09, CI 95% [0.04, 1.00]), (D) Speed (scalar) ($\chi 2$ Kruskal-Wallis (3) = 20.19, $p$ = 0.000155, effect size ($\varepsilon^2$) = 0.09, CI 95% [0.03, 1.00]). Statistical analysis was performed using Kruskal-Wallis non-parametric test with Dunn´s pairwise comparisons and Holm $p$ adjustment method (Supplementary table S13). For a better visualization, outliers are not shown in the figure, but highlighted in the Supplementary Excel file (Supplementary Table S12). Outlier analysis supports that *th* is required for normal kinematic performance.

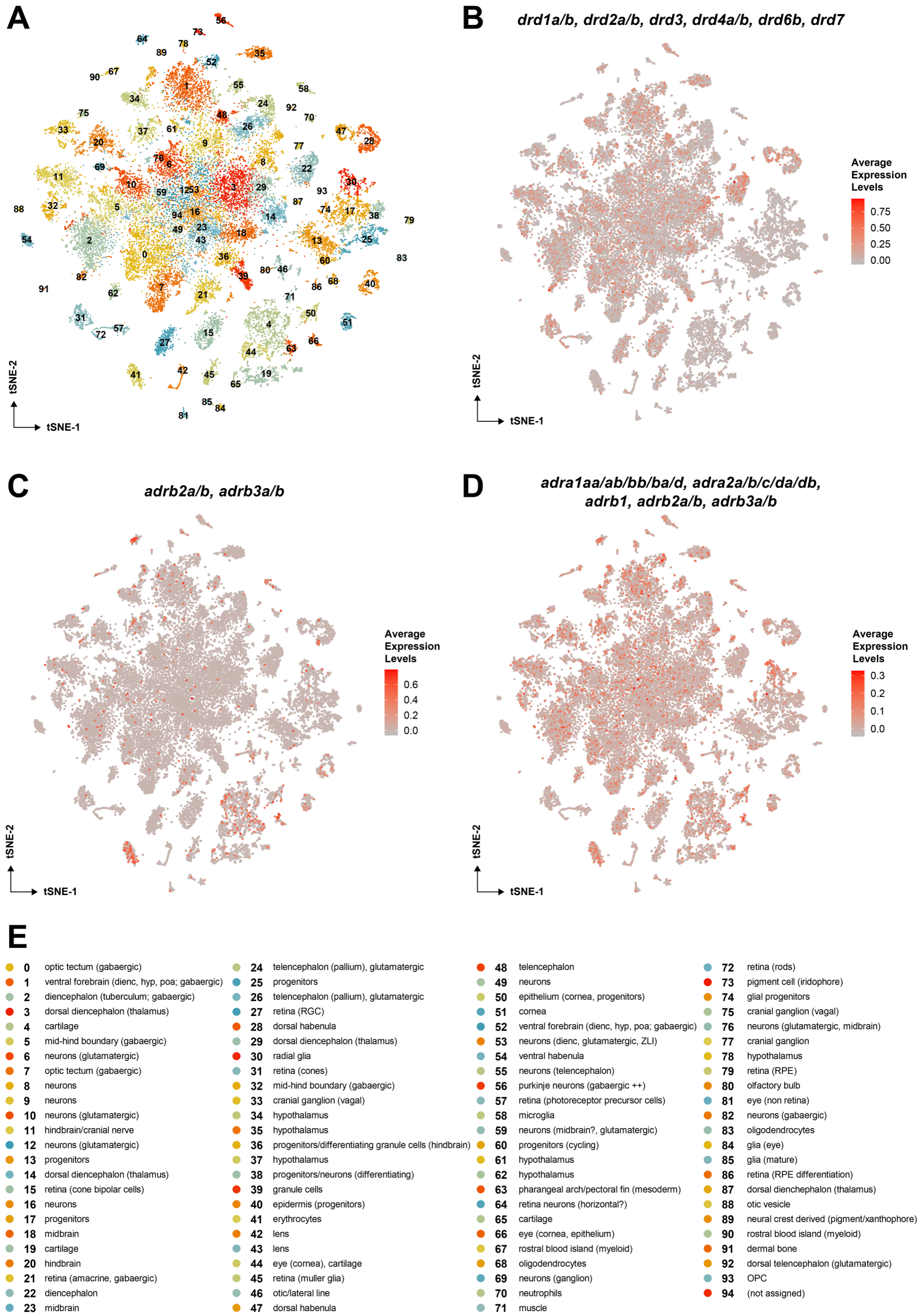


**Supplementary Figure S6. Expression of CA receptors in 5dpf brain cells**

scRNAseq Data from dissected brains and eyes from 5 dpf larval zebrafish published by Raj et al. (2020) were analyzed using the Seurat package (v4.3.0). **(A)** Color coded identified clusters from the publication, for annotation see (E). **(B-D)** Relative expression levels of CA receptors in single cells. **(B)** Expression of DA receptors, combined as indicated. **(C)** Beta2 and beta3-adrenergic receptors which have linked to noradrenaline signaling, combined as indicated. **(D)** Alpha and beta adrenergic receptors, combined as indicated. **(E)** Cluster Annotation was retained from original data.

**Supplementary Tables (EXCEL File)**

**Table S1: Figure 3 Dopamine ELISA**

Dopamine ELISA measurements for WT control dilution series and mutant genotypes. ANOVA analysis.

**Table S2: Figure 4C *th* cell counts**

Cell counts and statistical analysis of dopaminergic neurons for each analysed cell cluster in *th, th2* double mutants and WT siblings. For abbreviations of anatomical cluster names see Figure 4C legend.

**Table S3: Figure 5B cell counts 33 hpf**

Cell counts for the analysis shown in Figure 5B 33 hpf.

**Table S4: Figure 5B cell counts 3 dpf**

Cell counts for the analysis shown in Figure 5B 3 dpf.

**Table S5: Figure 6B HGC phenotypes 3 dpf**

Embryos from a *th+/-, th2+/-, tyr-/-* cross were analysed at 3 dpf for HGC phenotype. 180 embryos with HGC[+] phenotype were identified. 189 HGC[-] embryos were randomly selected from the cross as HGC[-] population. Genotypes for each embryo were determined by PCR after sorting.

**Table S6: Figure 6E levels of HGC retention among genotypes from 3 dpf to 5 dpf**

A random subset of 92 embryos classified as HGC+ retention phenotype at 3 dpf (Figure 6B and Table S5) were reanalysed at 5 dpf. The percentages were extrapolated based on the analysis group of 92 larvae with the 3 dpf HGC retention phenotype, and not on the total cross (see legend Table S5).

**Table S7: Figure 7 HR and HRV temperature dependence data**

HR and HRV measurements for the analysis shown in Figure 7 and in Supplementary Figure S3. Outliers are shown in red.

**Table S8: Figure 7 HR and HRV temperature dependence statistical analysis**

Two-way ANOVA and Tukey's multiple comparison results for HR and HRV related to Figure 7 and Supplementary Figure S3.

**Table S9: Figure 8A locomotor activity**

Spontaneous locomotor activity measurements of larvae obtained from *th+/-, th2+/-, tyr-/-* in-crosses. Analysis is shown in Figure 8A.

**Table S10: Figure 8A locomotor activity**

Spontaneous locomotor activity for control genotypes ABTL and *tyr-/-*. Analysis are shown in Supplementary Figure S4.

**Table S11: Figure 8C optomotor response**

Proportion of responders and non-responders among genotypes

**Table S12: Figure 9 optomotor response**

Raw values for trajectory, displacement and time were obtained with the manual tracking plugging from ImageJ and used to calculate directionality, speed and velocity. Since OMR was assessed in free swimming larvae, it was impossible to have the same initial position for all larvae, therefore it was not possible to make a comparison among raw variables.

**Table S13: Figure 9 optomotor response statistical tests**

Genotype distributions were compared using the two-sample Kolmogorov-Smirnov test. Pairs of compared genotypes are shown with an "x". D: Maximum distance between the empirical distribution functions of two samples. Results of Kruskal-Wallis with Dunn's pairwise comparisons and Holm *p* adjustment method for each kinematic parameter analyzed considering the outliers (Outliers Present; columns B-I) and not considering them (Outliers Removed; columns K-P).

**Supplementary Material References**

Altbürger, C., Holzhauser, J., & Driever, W. (2023). CRISPR/Cas9-based QF2 knock-in at the tyrosine hydroxylase (th) locus reveals novel th-expressing neuron populations in the zebrafish mid- and hindbrain. *Frontiers in Neuroanatomy, 17*. doi:10.3389/fnana.2023.1196868

Armbruster, D., Mueller, T., & Driever, W. (2025). Analysis of lhx8a, isl1, pax6a/b, calb2a and sst7 Reveals that Dopaminergic Neurons in the Zebrafish Subpallium Belong to the Extended Amygdala. *bioRxiv*, 2025.2001.2027.635026. doi:10.1101/2025.01.27.635026

Eisenhofer, G., Tian, H., Holmes, C., Matsunaga, J., Roffler-Tarlov, S., & Hearing, V. J. (2003). Tyrosinase: a developmentally specific major determinant of peripheral dopamine. *Faseb Journal, 17*(10), 1248-1255. doi:10.1096/fj.02-0736com

Filippi, A., Mahler, J., Schweitzer, J., & Driever, W. (2010). Expression of the paralogous tyrosine hydroxylase encoding genes th1 and th2 reveals the full complement of dopaminergic and noradrenergic neurons in zebrafish larval and juvenile brain. *Journal of Comparative Neurology, 518*(4), 423-438. doi:10.1002/cne.22213

Holzschuh, J., Ryu, S., Aberger, F., & Driever, W. (2001). Dopamine transporter expression distinguishes dopaminergic neurons from other catecholaminergic neurons in the developing zebrafish embryo. *Mech Dev, 101*(1-2), 237-243. doi:10.1016/s0925-4773(01)00287-8

Ma, P. M. (1994). Catecholaminergic Systems in the Zebrafish .1. Number, Morphology, and Histochemical-Characteristics of Neurons in the Locus-Coeruleus. *Journal of Comparative Neurology, 344*(2), 242-255. doi:DOI 10.1002/cne.903440206

Ma, P. M. (1997). Catecholaminergic systems in the zebrafish .3. Organization and projection pattern of medullary dopaminergic and noradrenergic neurons. *Journal of Comparative Neurology, 381*(4), 411-427.

Reiner, A., & Northcutt, R. G. (1987). An Immunohistochemical Study of the Telencephalon of the African Lungfish, Protopterus-Annectens. *Journal of Comparative Neurology, 256*(3), 463-481. doi:DOI 10.1002/cne.902560313

Rink, E., & Wullimann, M. F. (2001). The teleostean (zebrafish) dopaminergic system ascending to the subpallium (striatum) is located in the basal diencephalon (posterior tuberculum). *Brain Research, 889*(1-2), 316-330. doi:Doi 10.1016/S0006-8993(00)03174-7

Rink, E., & Wullimann, M. F. (2002). Development of the catecholaminergic system in the early zebrafish brain: an immunohistochemical study. *Brain Res Dev Brain Res, 137*(1), 89-100. doi:10.1016/s0165-3806(02)00354-1

Schredelseker, T., Veit, F., Dorsky, R. I., & Driever, W. (2020). Bsx Is Essential for Differentiation of Multiple Neuromodulatory Cell Populations in the Secondary Prosencephalon. *Front Neurosci, 14*, 525. doi:10.3389/fnins.2020.00525

Smeets, W. J. A. J., & Reiner, A. (1994). Catecholamines in the CNS of vertebrates: current concepts of evolution and functional significance. In W. J. A. J. Smeets & A. Reiner (Eds.), *Phylogeny and Development of Catecholamine Systems in the CNS of Vertebrates* (pp. 463-481). Cambridge: Cambridge University Press.

Yamamoto, K., Ruuskanen, J. O., Wullimann, M. F., & Vernier, P. (2010). Two tyrosine hydroxylase genes in vertebrates New dopaminergic territories revealed in the zebrafish brain. *Molecular and Cellular Neuroscience, 43*(4), 394-402. doi:10.1016/j.mcn.2010.01.006

Yamamoto, K., & Vernier, P. (2011). The evolution of dopamine systems in chordates. *Frontiers in Neuroanatomy, 5*. doi:10.3389/fnana.2011.00021
